# Genetic insights on the mechanisms of human cortical folding

**DOI:** 10.64898/2026.03.06.709690

**Authors:** William Snyder, Rebecca Shafee, Siyuan Liu, Elizabeth Levitis, Kuaikuai Duan, Kuldeep Kumar, Charles H Schleifer, Rune Boen, Christopher RK Ching, Joan C. Han, Nancy Lee, Jennifer G Mulle, Sarah Shultz, Sébastien Jacquemont, Carrie E Bearden, Petra E Vértes, Edward T Bullmore, Armin Raznahan

## Abstract

The unique and intricate pattern of human cortical folding is rooted in fetal neurodevelopmental processes and can now be comprehensively quantified by new neuroimaging-derived measures of sulcal complexity. Here, we provide the first genetic maps of human sulcal complexity. Beginning with large effects of rare variants, we survey nine different neurogenetic syndromes (*n*=615), detecting visible changes in sulcal complexity on a shared axis of sulcal change coupled to the prenatal timing of sulcation. Turning to common genetic variants, we use genome-wide association studies of complexity scores for 40 sulci in the UK Biobank (*n*∼29,000) to (i) resolve variable heritability across sulci, (ii) reveal both local and remote shared genetic effects with cortical morphology, and (iii) identify complexity-associated genes and their embedding in brain maps of prenatal gene expression. These reference genetic maps uncover multiple new mechanistic pathways for cortical morphogenesis in health and disease.

## Introduction

Cortical folding is one of the most visually striking features of human brain anatomy, with each individual’s brain having a unique folding pattern^1,2^. Individual folding geometry has important consequences for brain function, delineating functional boundaries in sensorimotor cortices^3,4^ and providing a scaffold upon which connectivity and communication between regions is constrained^5,6^. Because these folding patterns are established in prenatal life and remain stable across the life span^7,8^, studying postnatal or adult cortical folding patterns can provide a “rear-view mirror” reflecting much earlier brain developmental processes^7^.

The pronounced morphological complexity of human cortical folds (*i.e.*, sulci) offers a multitude of features which can be measured by neuroimaging and have been used to chart phenotypic variation in folding, its genesis, and its potential consequences. Thus, there have been numerous studies of interindividual variation across isolated sulcal features, such as sulcal depth^9–21^, depth variability^14,22–24^, length^12,13,19,21,25,26^, width^27–29^, orientation^2,30–33^, curvature^14,16,17,34,35^, and various gyrification indices^20,28,36–42^ – including evidence for disease and age-related sources of variation^5,40,43–49^. Some of these sulcal features have also been associated with both rare^50–53^ and common^21,27,54–60^ forms of genetic variation, hinting at a causal role for prenatal molecular programs, *e.g.* involving regulation of the progenitor pool for tangential expansion of the cortex^60–62^. However, existing studies of human cortical folding and its developmental mechanisms have yet to integrate these diverse separate features of sulcal morphology – despite growing evidence that these features are in fact highly coordinated with one another across prenatal development and across the cortical sheet^63^.

It has now become possible to combine multiple sulcal features from *in vivo* neuroimaging into a single integrative sulcal complexity score for each fold in an individual’s brain^63^. These scores are derived from an individual’s unique sulcal phenotype network (SPN), which represents the similarity between all pairs of sulci across multiple sulcal shape features. Although a given sulcus can vary greatly in its specific shape across individuals, the ranking of sulci by their relative complexity across the cortical sheet is highly stereotyped across individuals – spanning from consistently linear sulci (deep, variable depth, linear path; *e.g.*, central sulcus, calcarine fissure) to consistently complex ones (shallow, highly branched, fractal path; *e.g.*, orbitofrontal sulcus, medial parietal sulcus). Moreover, inter-fold differences in sulcal complexity are tightly correlated with when folds emerge in prenatal development (earliest for linear sulci, and latest for complex sulci)^63,64^, indicating that adult sulcal complexity provides a retrospective indicator of prenatal programs of cortical development. However, despite the capacity of sulcal complexity scores to efficiently summarize multiple morphological features of human cortical folds, and track differences in the timing of their prenatal emergence, nothing is yet known regarding the genetic architecture of sulcal complexity. Addressing this gap in knowledge would provide a major step forward in clarifying the mechanisms that shape early human brain development.

Here, we provide precise mapping of genetic effects on human sulcal complexity, drawing from rare genetic variation in neurogenetic syndromes and from common genetic variation in a population-level imaging genomics cohort. We predicted that these complementary samples would reveal separable pathways of genetic influence on cortical folding, either by disrupting canalized neurodevelopmental programs for sulcation (rare variation) or by exerting distributed, pleiotropic influences on the developing cortical sheet (common variation). First, we collated structural brain magnetic resonance imaging (MRI) from nine neurogenetic syndrome cohorts with a chromosomal aneuploidy or copy number variation (CNV), which we refer to as follows (including dosage nomenclature used in figures and alternative names): XXY (+X, Klinefelter syndrome), XYY (+Y), 3q29 deletion syndrome (−3q29), 11p13 deletion syndrome (−11p13, WAGR syndrome), 16p11.2 deletion syndrome (−16p11.2), 16p11.2 duplication syndrome (+16p11.2), trisomy 21 (+21, Down syndrome), 22q11.2 deletion syndrome (−22q11.2, Velocardiofacial Syndrome), and 22q11.2 duplication syndrome (+22q11.2). These recurrent gene dosage disorders all have well-established impacts on neurodevelopment and behavior, some distinct and some shared^65–67^. Thus, this uniquely large and diverse MRI dataset allowed us to test whether such rare genetic variation also places either distinct or shared systematic burdens on programs for cortical folding – such as by disrupting the temporal window for fetal sulcal emergence that mediates linear-to-complex sulcal shape differentiation.

To identify common variant effects on sulcal complexity, we leveraged the large sample of the UK Biobank with both structural brain MRI and genomics (*n*∼29,000). We performed comprehensive annotation of common variant genetic influences on sulcal complexity by: estimating regional variation in the SNP-based heritability of sulcal complexity; examining the shared genetic architecture of sulcal complexity with mechanistically linked parameters of the cortical sheet; characterizing individual variants and genes with significant GWAS signal for sulcal complexity; and, contextualizing these genes in transcriptional gradients of the fetal brain. Together, our findings delineate multiple routes for the genetic regulation of sulcal complexity, providing fresh insights into the mechanistic basis of cortical folding.

## Results

### Sulcal phenotype networks (SPNs)

Sulcal complexity was computed as in Snyder et al.^63^ Briefly, five sulcal phenotypes were measured for each of 40 sulci identifiable in brain MRI (**Fig. 1A**, **Extended Data Fig. 1**): average depth (AD), depth variability (DV), longest branch (LB), branch span (BS), and fractal dimension (FD); spanning radial (AD, DV) and tangential (LB, BS, FD) dimensions of cortical folding. Pairwise correlations of these five features between any two sulci defined individual SPNs, providing a fingerprint of shape similarities within a cortex. As the first principal component of a group mean SPN captures the linear-to-complex patterning across sulci^63^, sulcal complexity was derived in all analyses as the coherence of a subject’s sulcus (individual SPN row) with the first principal component of their matched controls’ mean SPN (**Fig. 1A**, **Supplementary Fig. 1**, **Methods**) – forming a single composite metric scoring all sulci along a linear (deep, variable depth, linear path) to complex (shallow, fractal) shape axis.

**Figure 1.**
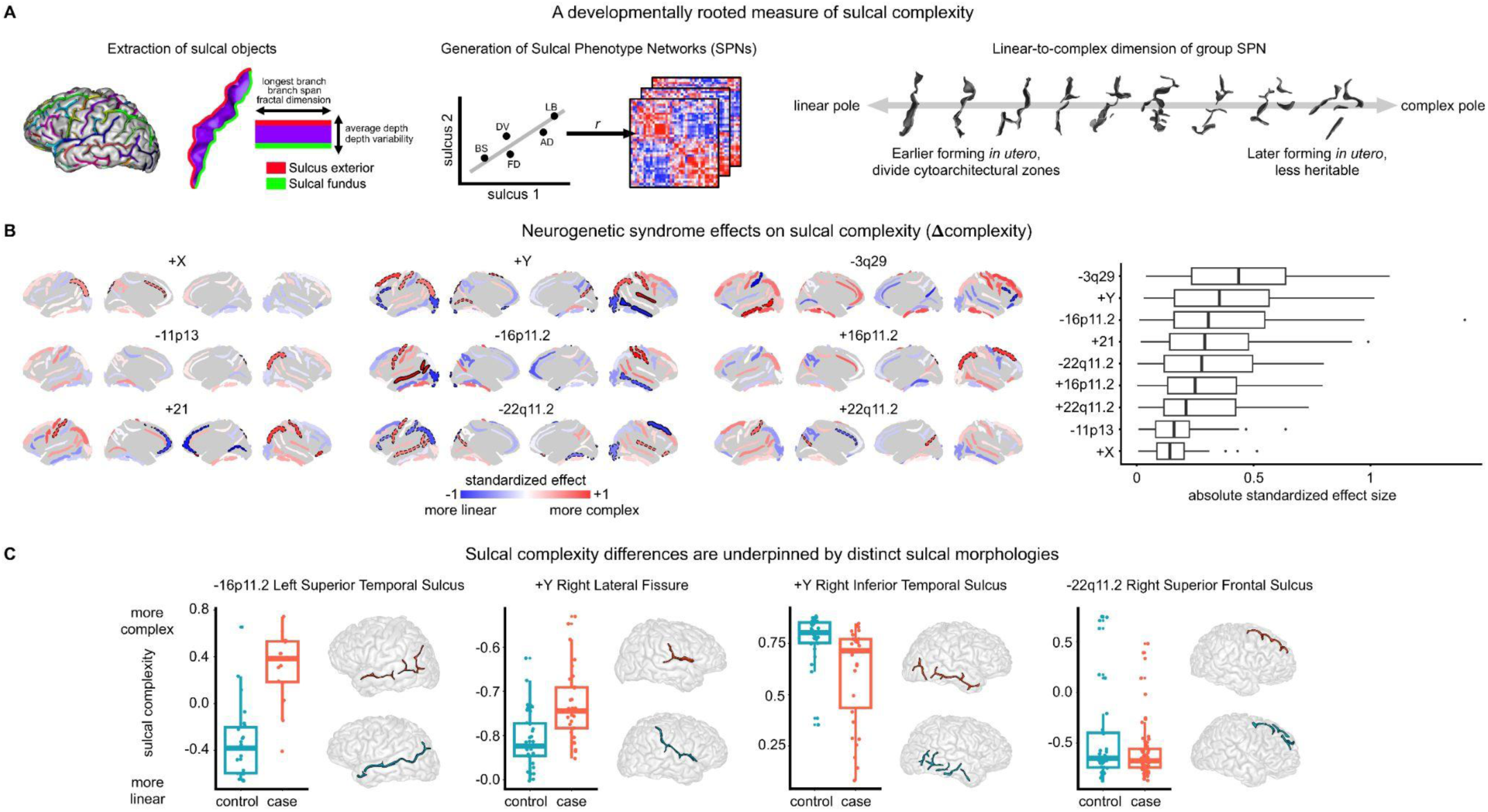
Sulcal phenotype networks have anatomically diverse patterns of atypical complexity across neurogenetic syndromes. **A)** For all analyses, sulcal objects are extracted as the space between gyri for 40 sulcal regions, enabling morphological measurement of 5 sulcal phenotypes per sulcus for individual sulcal phenotype network (SPN) computation. Sulcal complexity is derived from SPNs as the coherence of a subject’s fold (SPN row) with the first principal component of matched controls’ mean SPN – which consistently ranks folds along a linear-to-complex shape axis (**Supplementary** Fig. 1). **B)** The standardized effect sizes for each of nine different neurogenetic syndromes were computed for the complexity scores of each of 40 different sulci. Positive (red) effects indicate the sulcus is more complex in cases than controls, and negative (blue) effects indicate sulci that are more linear in cases. Bold sulci are significant following Bonferroni correction across sulci within syndromes, dash-outlined sulci are nominally significant. All effects and exact *p*-values are provided in **Supplementary Data 1.** In the boxplots, syndromes are ordered by rank median absolute standardized effect size. Boxplots on the right panel show the distribution of sulcal complexity effect sizes across the 40 sulci for each of the nine neurogenetic syndromes. **C)** Raw sulcal complexity scores are plotted for each of the four (Bonferroni-corrected) significant sulci, ordered left-to-right by decreasing standardized absolute effect size. Raw sulcal complexity scores are shown to highlight the interpretability of its values even prior to fitting to a linear model; *i.e.*, sulci like the left superior temporal sulcus can be reliably interpreted as complex with an interruption in its branches when uncorrected sulcal complexity > 0. Source Data are provided as Source Data files.

### Stability of neurotypical sulcal architecture

The linear-to-complex patterning of cortical folding across cortical regions is highly constrained in neurotypical adults^63^ but has not yet been investigated in younger populations. Therefore, before analyzing sulcal complexity in youth with neurogenetic syndromes, we first evaluated whether sulcal architecture was conserved in the age-matched control groups for each clinical cohort (combined control *n*=312, comprised of one control group for each of nine neurogenetic syndrome cohorts, **Supplementary Tables 1-2**). Mean SPNs for each control group were almost identical to each other (mean edge-wise correlation *r* = 0.98, cohorts’ mean age range: 13-37 years) and to the group mean SPN first described in the older UK Biobank cohort (mean *r* = 0.95, UK Biobank age range: 45–82 years) (**Extended Data Fig. 2**). This indicates the high replicability of sulcal patterning represented by SPNs in multiple independent cohorts and suggests that the linear-to-complex axis of cortical folding is unchanging over the course of normal adult life.

### Rare genetic variant effects on sulcal complexity

As an initial probe for genetic influences on human sulcal complexity, we focused on nine different rare neurogenetic syndromes (total case *n*=313, control *n*=312, **Supplementary Tables 1-2**) with known large effects on neurodevelopment^65–75^. The diverse genetic bases for these nine conditions – spanning aneuploidies and sub-chromosomal copy number variations – provided a powerful test for convergent vs. specific influences of rare genetic variation across measures of sulcal complexity at each of 40 sulci.

The nine neurogenetic syndromes studies had anatomically distinct patterns of atypical sulcal complexity (Δcomplexity) compared to matched controls, as measured using linear models for each sulcus (**Fig. 1B**, **Methods**). Individual sulcal complexity scores were not sensitive to computation using any particular cohort’s mean control group SPN (**Supplementary Fig. 1**), reinforcing the reproducibility of control SPNs (**Extended Data Fig. 2**), and facilitating Δcomplexity comparisons between syndromes.

Across the nine neurogenetic syndromes, four sulci showed statistically significant differences from controls after correction for multiple comparisons across sulci within each syndrome (*p* < 0.05 / 40 sulci), and 49 out of all 360 tested sulci showed nominally significant (*p* < 0.05) case-control differences in sulcal complexity of moderate-to-large absolute standardized effect size (0.43-1.40). Across syndromes, convergent, strong effects (|Δcomplexity| > 0.4) were observed for the left post-central sulcus, right central sulcus, and bilateral intra-parietal fissures (**Fig. 1B**, **left panel**). The distribution of Δcomplexity values varied across syndromes (**Fig. 1B**, **left panel**), with 3q29 deletion syndrome conferring the strongest cortex-wide alterations in sulcal complexity (mean absolute standardized effect size = 0.45) while XXY exhibited the most modest effects (mean absolute standardized effect size = 0.15). These neurogenetic effects on sulcal complexity were estimated by a linear model including brain total tissue volume (TTV) as a covariate but were replicable when not controlling for TTV (**Supplementary Fig. 2**), suggesting sulcal complexity changes in neurogenetic syndromes index robust size-independent changes in sulcal shape.

Atypical sulcal complexity detected by SPN analysis was linked to visible differences in folding of specific sulci in specific syndromes (**Fig. 1C**). Notably, individual sulcal complexity scores > 0 in the left superior temporal sulcus detected an interruption in the continuity of this sulcus, a landmark almost exclusively associated with 16p11.2 deletion cases relative to their matched controls. Visually salient differences were also observed in other syndromes, such as more Y-shaped right lateral fissure in XYY, less interrupted branching in XYY right inferior temporal sulcus, and more linear path of the right superior frontal sulcus in 22q11.2 deletion syndrome. These distinctive morphologies suggest atypical mechanisms in the fetal emergence of the sulcal roots that arrange to form larger sulcal regions^22,76^, imprinting lasting traits of atypical interruptions or orientations of these roots within sulci.

### Comprehensive phenotypic profiling of sulcal alterations in rare genetic disorders

To interrogate the morphological accompaniments of sulcal complexity changes, we: (i) investigated a battery of 7 other sulcal phenotype effects in each neurogenetic syndrome (**Extended Data Fig. 3A-B**, **Supplementary Data 1**), and (ii) quantified the cross-sulcus spatial congruence between changes in these 7 specific sulcal metrics and changes in sulcal complexity for each neurogenetic syndrome (**Methods, Extended Data Fig. 3C**). These analyses revealed that changes in average depth and branch span were most strongly linked to changes in sulcal complexity (median spatial correlation *r* = −0.46, 0.61, respectively) (**Extended Data Fig. 3C**). These two metrics span radial and tangential orientations of the cortical sheet, respectively, highlighting coordination of altered sulcal development along multiple axes of cortical folding. Changes in sulcal surface area and sulcal thickness, frequently studied in neurogenetic syndrome populations^68–70,72,73,75^, were strikingly not spatially correlated across sulci with changes in sulcal complexity – despite being parameters thought to mechanistically influence degrees of cortical folding^77–79^ (**Extended Data Fig. 3C**). We therefore demonstrate that sulcal complexity offers a unique signal to probe genetic effects on early life cortical development.

### A shared spatial axis of atypical sulcal complexity in rare genetic disorders

Since the neurogenetic syndromes investigated here converge on early-life behavioral presentation of developmental or psychiatric disorder, we hypothesized that (1) these diverse neurogenetic syndromes may also show spatial convergence in their effects on sulcal complexity, and (2) spatial axes of shared atypical cortical folding would be linked to known spatial axes that govern fetal sulcal development or sulcation. Pairwise spatial correlations between neurogenetic syndrome Δcomplexity effect maps (**Fig. 2A**) demonstrated novel within- and between-syndrome patterning of cortical vulnerability and did indeed reveal shared spatial axes of sulcal complexity effects (Δcomplexity) across the nine diverse neurogenetic syndromes studied (**Fig. 2B-C**).

**Figure 2.**
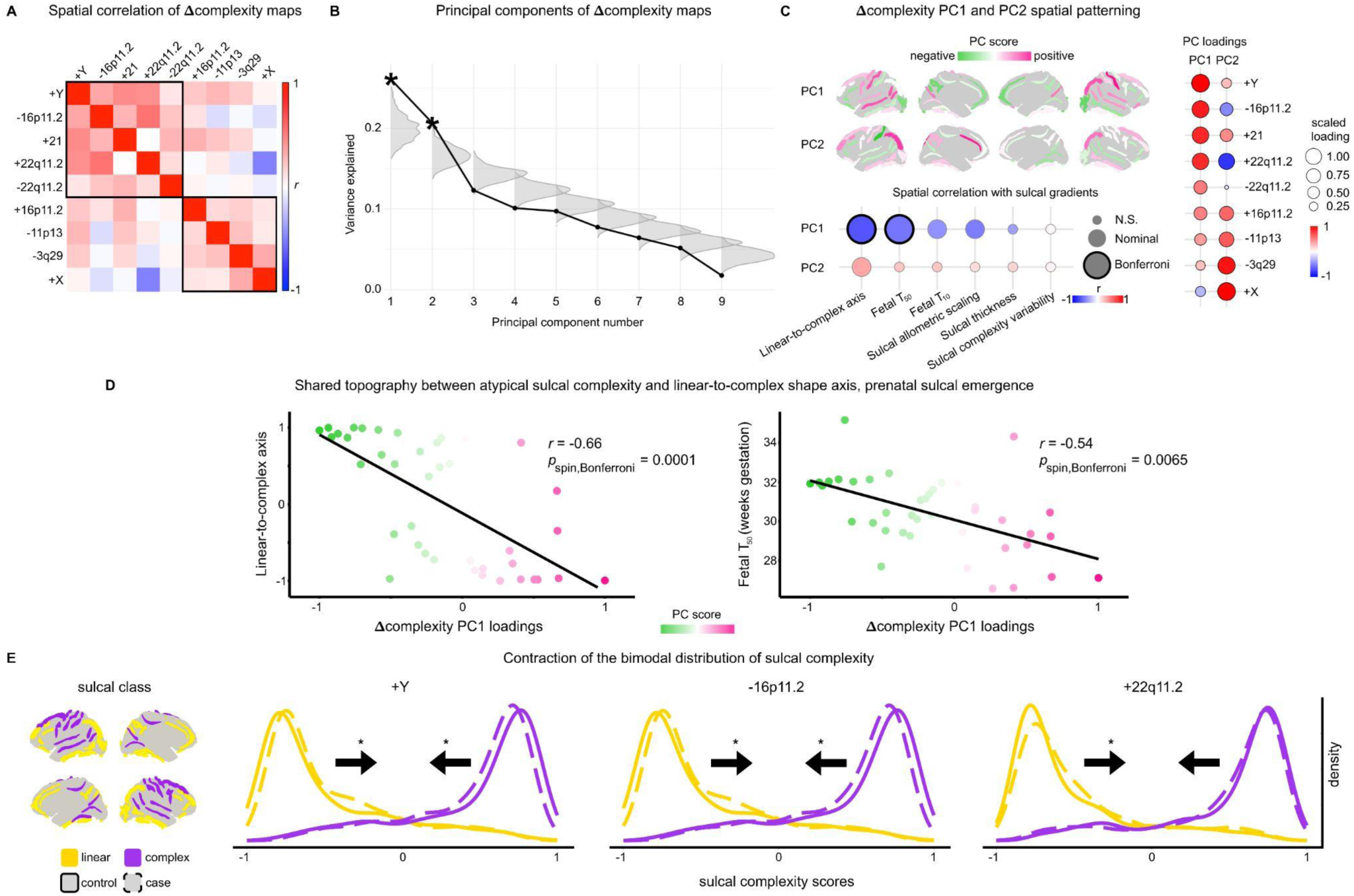
Convergent sulcal complexity changes across neurogenetic syndromes are spatially coupled to the timing of prenatal sulcal emergence. **A)** Spatial correlations between maps of atypical sulcal complexity in neurogenetic syndromes (Δcomplexity) are shown, with syndromes in the correlation plot rank-ordered by the first principal component loadings of cross-syndrome variability in spatial patterning. **B)** The first two PCs of atypical sulcal complexity – collectively explaining 47% variance across neurogenetic syndromes – explain significantly more variance in the spatial distribution of atypical sulcal complexity effects across neurogenetic syndromes than randomly permuted spatial maps (gray distributions) (*p*_perm_ < 0.05). **C)** The first two PCs of atypical sulcal complexity are plotted on the cortical surface (Δcomplexity PC1, PC2), with the sign of PC scores chosen to be positively correlated with the syndrome Δcomplexity effects that most strongly load on respective PCs. Syndrome loadings onto PCs are scaled to [-1,1] for comparison between PCs. These dominant modes of rare genetic variant effects were annotated for their spatial coherence with gradients of sulcal morphology. **D)** Significant spatial correlations with Δcomplexity PC1 are plotted, demonstrating shared topography between the dominant mode of rare genetic variant effects and (1) the linear-to-complex patterning of typical sulcal shape across the cortical sheet and (2) the timing of when each sulcus reaches 50% folded *in utero* (Fetal *T*_50_). PC1 scores are scaled to [-1,1]; positive values are observed in typically linear sulcal regions (*e.g*., central sulcus) and negative values are observed in typically complex sulcal regions (*e.g.*, occipital sulcus), indicating a contraction of the linear-to-complex axis linked to fetal sulcal emergence. **_E)_** Contraction of the linear-to-complex axis is directly demonstrated for XYY, 16p11.2 deletion, and 22q11.2 duplication syndromes, with shifting of distributions away from extreme linear or complex poles of morphology. Sulci were grouped into linear or complex classes based on their average neurotypical complexity scores^63^ (left). Asterisks denote significant shifts of distributions for highly linear or highly complex sulcal classes (*p*_linear_ or *p*_complex_ < 0.05). Note that the effect of 22q11.2 duplication syndrome on complexity scores for the complex sulcal class resulted in negative shifts consistent with contraction but not reaching nominal significance. Raw sulcal complexity scores are shown; however, covariate residualized scores were used in linear models comparing cases and controls (**Methods**). This contraction or dedifferentiation of sulcal complexity in syndromes could reflect contractions of the prenatal window for sulcal emergence that reinforces extremes of linear vs. complex sulcal shape. Source Data are provided as Source Data files.

The shared topography of atypical sulcal complexity between syndromes deviated from previously described morphometric changes in two notable regards. First, in contrast to the mirrored volume, surface area, or thickness changes seen between individuals with duplications versus deletions of the same locus^68,69,71,74,75^, we observed weak positive correlations between syndromes paired by CNV dosage (+/-16p11.2, +/-22q11.2; *r* = 0.11, 0.14, respectively). Rather, both increased and decreased dosage of genes in these CNVs have capacity to convergently change sulcal architecture. Second, XXY and XYY sulcal complexity effects were effectively uncorrelated (*r* = 0.04), in contrast to their strongly coupled regional effects of XXY and XYY syndrome on cortical volume, surface area, and thickness^73,80^. These two observations further underline the distinct information provided by mapping genetic effects in sulcal complexity vs. cortical volume, area and thickness.

Spatial comparison of Δcomplexity maps between syndromes revealed two separate spatial modes of atypical sulcal complexity, with moderate cross-syndrome spatial correlation within each of these two syndrome clusters (**Fig. 2A**, top-left cluster median *r* = 0.29, bottom right *r* = 0.16). These two distinct spatial axes of sulcal complexity changes were also evident from principal component analysis (**Methods**), which detected two significant (*p*_perm_ = 0.002, 0.001, respectively) principal components (Δcomplexity PC1 and PC2, **Fig. 2B-C**, **Supplementary Data 2**) that recapitulated the two syndrome clusters (**Fig. 2A**).

To aid interpretation of these newly discovered convergent spatial axes of rare variant effects on sulcal complexity, we quantified their alignment with known gradients of sulcal morphology and development. The dominant mode of rare genetic variation effects – Δcomplexity PC1 – most strongly spatially cohered with the linear-to-complex regional organization of cortical folding (*r* = −0.66, *p*_spin_,_Bonferroni_ = 0.0001) and the fetal gestational age when a sulcus reaches 50% of its final sulcation (Fetal *T*_50_; *r* = −0.54, *p*_spin_ = 0.0065) (**Fig. 2C-D**). Thus, Δcomplexity PC1 represents a contraction of the linear-to-complex sulcal axis of the human brain: earlier-forming, linear sulci become more complex and complex sulci become more linear in response to rare genetic variation. This phenomenon was associated most with XYY, 16p11.2 deletion, trisomy 21, and 22q11.2 deletion and duplication syndromes (**Fig. 2C**, PC loadings). The strongest direct evidence for contraction or de-differentiation of complexity scores – *i.e.*, positive shifts of complexity scores for consistently linear sulci and negative shifts for consistently complex sulci (**Methods**) – was observed for XYY (*p_linear_* = 0.00081, *p_complex_* = 0.0030), 16p11.2 deletion (*p_linear_* = 0.015, *p_complex_* = 0.044), and 22q11.2 duplication syndromes (*p_linear_* = 0.013, *p_complex_* = 0.076) (**Fig. 2E**).

Δcomplexity PC2 nominally exhibited an expansion of linear-to-complex organization (*r* = 0.34, *p*_spin_ = 0.049), while also highlighting regions that are most consistently atypical across syndromes (*e.g.*, left post-central sulcus, left and right intra-parietal fissure). Δcomplexity PC2 was associated most with XXY, 3q29 deletion, 11p13 deletion, and 16p11.2 duplication syndromes (**Fig. 2C**, PC loadings). Both Δcomplexity PC1 and PC2 spatial axes were replicable when not controlling for effects of TTV (**Supplementary Fig. 3**).

Taken together, these analyses show that multiple diverse rare genetic variants can induce sulcal complexity changes that are orthogonal to previously studied aspects of cortical anatomy, and, in some cases, result in visible changes that can distinguish cases and controls. Moreover, we find that the effects of nine diverse rare genetic variants on cortical folding converge on common spatial axes of atypical sulcal complexity that are tied to the sequential emergence of cortical folds in prenatal life – hinting at a close link between the temporal and spatial patterning of cortical folding variation, contributing to contractions or expansions of expressed sulcal complexity.

### Common variant (SNP-based) heritability of sulcal complexity

Given the substantial effects of rare genetic variants on sulcal complexity (**Figs. 1-2**), we next tested if and how sulcal complexity is influenced by common genetic variation. Previous sulcal GWAS have investigated a variety of individual sulcal measures^21,27,54–60^; but, to date, none have integrated multiple morphometrics. We therefore used the UK Biobank dataset (*n*∼29,000; mean age = 64 years; 46% male) to perform a GWAS of sulcal complexity (**Methods**) for the same set of 40 folds studied in the neurogenetic syndrome cohorts.

We first applied Linkage Disequilibrium Score Regression (LDSC)^81^ to GWAS summary statistics (i) to estimate the SNP-based heritability of sulcal complexity for each fold (**Fig. 3A**, left panel), and (ii) to compare the topography of complexity heritability with other spatial gradients of sulcal complexity in typical development and in rare (neurogenetic atypical) effects on sulcal complexity (**Fig. 3A**, right panel). The magnitude of sulcal complexity heritability varied considerably across the cortex (**Fig. 3A**, **Supplementary Data 3**); and this topographic patterning of sulcal heritability was highly replicated in a sensitivity analysis controlled for genetic effects on TTV (*r* = 0.99, **Supplementary Fig. 4**). 25 of 40 sulci had significant heritability (*p*_Bonferroni_ < 0.05, *h*^2^ range = [0.05, 0.1], median = 0.07) and 37 out of 40 sulci reached nominal significance (*p* < 0.05, *h*^2^ range = [0.06, 0.1], median = 0.06). Having resolved the cortical topography of SNP-based heritability for sulcal complexity, we next compared it to several key sulcal complexity gradients in health, as well as the dimensions of atypical sulcal complexity from rare genetic variation (**Fig. 2C**). The direction of spatial correlations followed the expected trend that more heritable sulci should be the most linear and earlier-emerging (**Fig. 3A**, right panel: linear-to-complex axis, *T*_10_, and *T*_50_ all approximately *r* = −0.29). However, these associations and associations with rare neurogenetic syndrome atypical sulcal complexity were not statistically significant even for more moderate correlations (|*r*| ∼ 0.3, *p*_spin_ > 0.05), potentially due to this pattern being driven by or most detectable from subsets of all cortical sulci. Notably, three sulci that were among the most repeatedly impacted across rare neurogenetic syndromes (average absolute standardized effect = 0.40, average percentile = 0.92) – the left and right intra-parietal fissures and the left superior temporal sulcus – were also amongst those showing the highest SNP-based heritability (average *h*^2^ = 0.09, average percentile = 0.90) – showing that the sulci which were most consistently atypical in neurogenetic syndromes were also more heritable in the general population.

**Figure 3.**
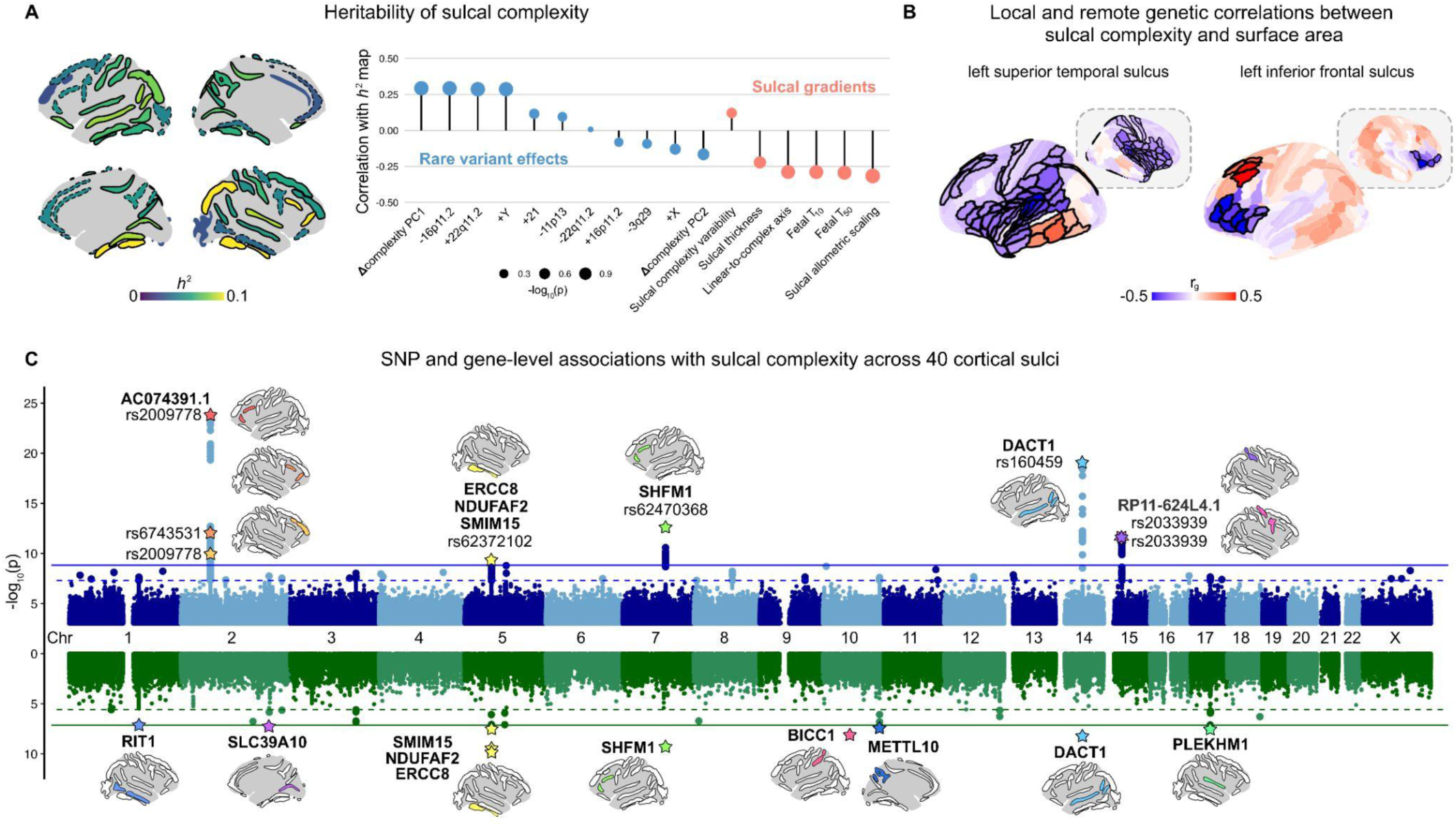
The common variant architecture of sulcal complexity is shared with local and remote cortical anatomy and maps to specific genomic regions. **A)** SNP-based heritability of sulcal complexity varies across sulci (significant SNP-based heritability is shown with bolded outlines *p*_Bonferroni_ < 0.05 and nominally significant sulci shown with dashed outline *p* < 0.05). Strongest heritability is observed for three sulci on the right hemisphere: the right intra-parietal fissure, right occipital-temporal lateral sulcus, and right inferior frontal sulcus (all *h*^2^ = 0.10). The three sulci not reaching nominal significance for heritability are all sulci with high average complexity scores: the left intermediate frontal sulcus, left anterior cingulate sulcus, and right occipital sulcus. Spatial correlations with the sulcal complexity *h*^2^ map do not reach statistical significance for rare genetic variant effects (atypical sulcal complexity, Δcomplexity) or sulcal gradients defined in normative population, but are signed as predicted; positive correlations indicate that more heritable sulci are more likely to emerge earlier *in utero* and are on average more linear; positive correlations with Δcomplexity PC1 indicate that more heritable sulci (which tend to be more linear) are more likely to have increased complexity in neurogenetic syndromes. **B)** Genetic correlations between sulcal complexity and cortical surface area exhibit spatially distinct associations surrounding and remote to each sulcus. Significant positive and negative associations are bolded and corrected across cortical parcels (FDR *q* < 0.05). For the two exemplar sulci shown, positive and negative genetic correlations flank the gyri adjacent to where the named sulcus would lie, meaning that the common genetic variation that predicts increased sulcal complexity would predict a pattern of both increased and decreased surface area around either side of the sulcus. Notably, even distal or remote regions and left-right homologous regions (gray dashed-outline boxes) exhibited significant genetic correlations with sulcal complexity for an individual sulcus. Additional exemplar sulci displaying this pattern of local and remote genetic correlations with area are shown in **Extended Data** Fig. 4A. Additional spatial patterns were observed for other sulci and also with cortical thickness, demonstrating both local and global (*i.e.*, cortex-wide) genetic associations with sulcal complexity (**Extended Data** Fig. 4A**-B**, **Supplementary** Figs. 5 and 6). **C)** SNP- and gene-level associations with sulcal complexity across 40 cortical sulci. Manhattan plots display the SNP (top) and gene (bottom) associations with sulcal complexity, with all associations for all 40 sulci’s GWAS overlaid. Solid lines indicate experiment-wide significance (*p*_SNP_ < 1.47 ×10^−9^, *p*_gene_ < 7.6 ×10^−8^) and dashed lines indicate genome-wide significance (*p*_SNP_ < 5 ×10^−8^, *p*_gene_ < 2.6 ×10^−6^). For SNP associations (top), lead SNPs for experiment-wide significant loci for each sulcus are labeled (stars), with gene (or lincRNA) names additionally labeled where loci fall within 10kb of the gene (or lincRNA). For gene associations (bottom), p-values from MAGMA^82^ are displayed, with experiment-wide significant genes labeled (stars). Source Data are provided as Source Data files.

### Genetic correlations between sulcal complexity and other aspects of cortical anatomy and neurodevelopment

We next used GWAS summary statistics for regional sulcal complexity to test for evidence of shared genetic architecture between sulcal morphology and other salient neuroanatomical and neurobehavioral phenotypes. We began by determining genetic correlations between sulcal complexity and regional measures of cortical surface area and thickness (**Methods**). These two morphometric features: capture the two sole defining axes of the cortical sheet that together determine cortical volume^83–85^; represent major foci of prior neuroimaging research^15,66,70,86–89^; and have both been closely linked to the biomechanics of tissue propensity to fold^77,78^. We found that sulcal complexity shows both local and remote genetic relationships with cortical surface area and thickness.

For many sulci, sulcal complexity had significant positive and negative genetic correlations with the surface area of cortical regions directly on either side of the fold (FDR corrected across 360 cortical parcels, *q* < 0.05; **Fig. 3B**, **Extended Data Fig. 4A**, **Supplementary Fig. 5**, **Supplementary Data 4**). For example, genetic factors imparting greater complexity of the left superior temporal sulcus tend to increase the surface area of immediately ventral lateral temporal cortices and decrease the surface area of immediately dorsal insula and frontal cortices (**Fig. 3B**). A similar effect was seen for the right paracingulate, left inferior frontal and right post-central sulci (**Fig. 3B**). Several folds also showed local genetic correlations with cortical thickness (**Extended Data Fig. 4B**, **Supplementary Fig. 6**). These patterns of local genetic correlation newly buttress previously proposed models for the genesis of cortical folding in which trans-sulcal genetic gradients defining arealization of the developing cortex contribute to sulcal formation^4,63,90,91^. Our findings also extend these models, by revealing that genetic factors increasing the complexity of a given fold can also regulate the surface area of specific cortical regions that are remote from the fold. This is seen, for example, between the left superior temporal and inferior frontal sulci and regional surface area surrounding their contralateral homologs. Remote genetic correlations are also apparent ipsilaterally – as seen between the left superior temporal sulcus and surface area of the left dorsolateral prefrontal cortex (**Fig. 3B**).

We additionally estimated genetic correlations with various psychiatric disorders and developmental phenotypes (**Extended Data Fig. 5**), interestingly identifying only weak or insignificant associations – in contrast to the above strong surface area and thickness genetic correlations. These findings show that while common genetic influences on sulcal complexity are largely uncoupled from those on behavioral traits (as is the case for most other regional neuroanatomical phenotypes^21,55,60^), they are intimately linked with genetic influences on cortical surface area and thickness.

### Genomic loci influencing sulcal complexity

To chart specific sources of genetic control over sulcal complexity, we next determined SNP- and gene-level associations for each of 40 sulci across the cortex (**Fig. 3C**). At the SNP-level, we identified 5 independent loci associated with sulcal complexity, linked to 8 different sulci (*p*_SNP_ < 1.47×10^−9^; **Fig. 3C**, top plot). Three loci—associated with the right occipital-temporal lateral sulcus, left inferior frontal sulcus, and left superior temporal sulcus—all mapped to genes proximal to or containing the loci (ERCC8, NDUFAF2, SMIM15, SHFM1, DACT1), as identified by MAGMA^82^. The other two loci were in intergenic regions. On chromosome 2, the left inferior frontal sulcus, right inferior frontal sulcus, and right intermediate sulcus—all typically complex sulci in the frontal cortex—had significant loci in the long intergenic non-coding RNA (lincRNA) AC074391.1 (also termed LINC02934) (**Extended Data Fig. 6**). Similarly, on chromosome 15, the right pre-central sulcus and right post-central sulcus mapped to lincRNA RP11-624L4.1 (LINC02915) (**Extended Data Fig. 6**). Congruent with our findings of local genetic correlations between sulcal complexity and cortical area and thickness (**Fig. 3B**), we found that for several folds, SNPs associated with sulcal complexity in the current study had shown genome-wide significance for measures of cortical area and thickness at regions surrounding the fold in prior work using an independent cortical parcellation (**Extended Data Figure 4B**).

Using MAGMA, we identified 10 genes associated with regional sulcal complexity at experiment-wide significance (*p*_gene_ < 7.6 × 10^−8^). 5 of the experiment-wide significant genes (ERCC8, NDUFAF2, SMIM15, SHFM1, DACT1) were associated with aforementioned experiment-wide significant SNP loci while the other 5 represented composite effects of SNPs in the genes’ locus (RIT1, SLC39A10, BICC1, METTL10, PLEKHM1). 33 candidate sulcal complexity genes exhibited genome-wide significance with MAGMA (*p*_gene_ < 2.6 ×10^−6^; Fig 3C, bottom plot), in addition to 17 genes identified as being within 10kb of genome-wide significant SNPs (*p*_SNP_ < 5 ×10^−8^), for a total of 50 candidate sulcal complexity genes (**Extended Data Fig. 7**). Many of these 50 genes are annotated as being involved in core aspects of fetal development (*e.g.*, regulating early transcription, differentiation and morphogenesis) and with certain mutations associated with neurodevelopmental disorders or cortical malformations – suggesting biological mechanisms for their influence on cortical folding (**Supplementary Data 5**). As a set, these 50 genes did not show statistically significant enrichment for any specific biological processes or cellular compartment gene ontology categories.

### Expression of sulcal complexity genes in the prenatal cortex

Given the prenatal timing of cortical fold emergence, we sought to contextualize genes that causally influence sulcal complexity (**Fig. 3C**) against gradients of gene expression in the developing fetal cortex when folds first emerge. To achieve this, we harnessed the µBrain dataset^58^, which provides measures of gene expression from 20 cortical regions and 5 tissue layers from the fetal human brain at both 16 and 21 post-conception weeks (pcw) (**Fig. 4A, Methods**).

**Figure 4.**
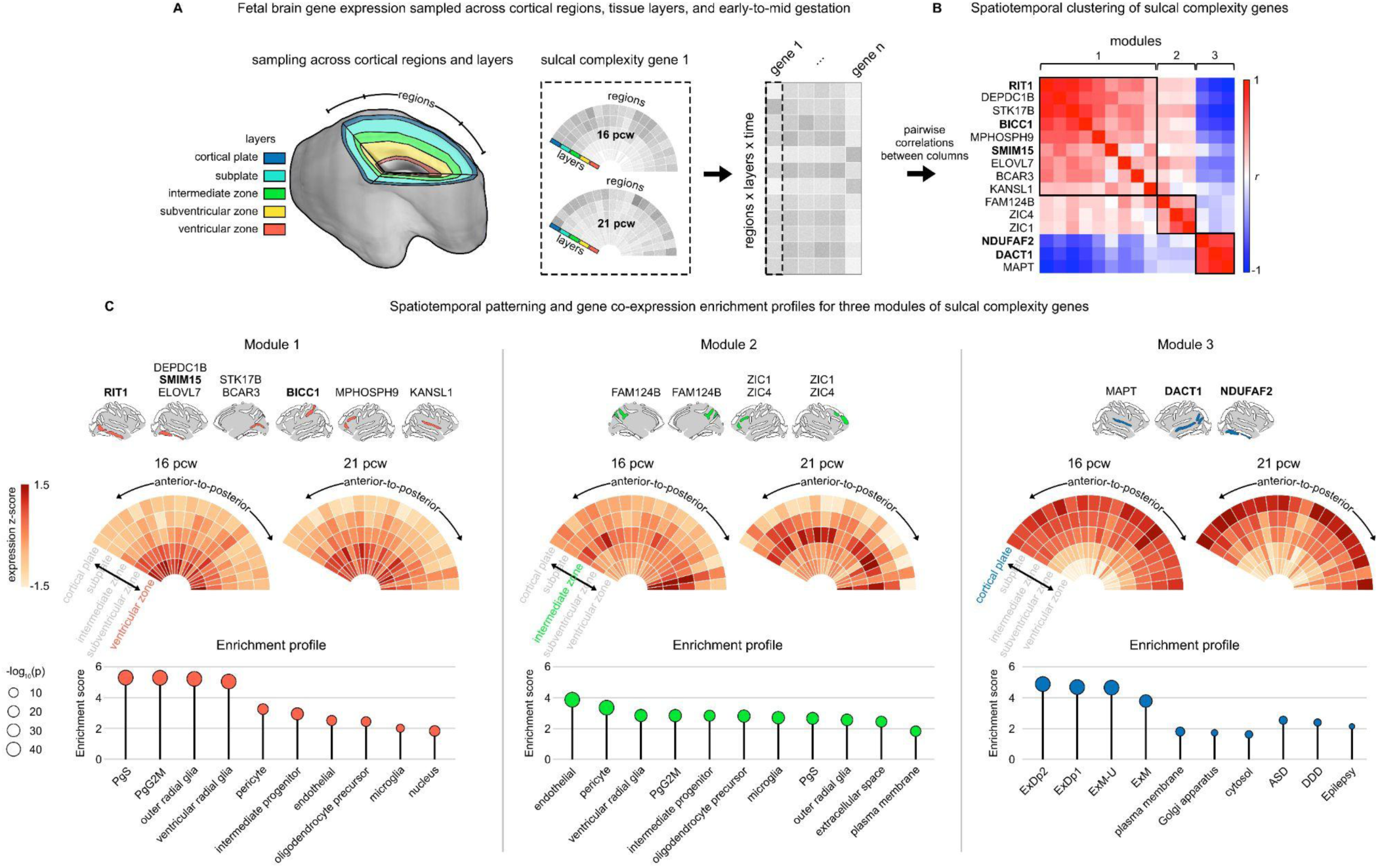
Sulcal complexity genes comprise three modes of fetal spatiotemporal gene expression associated with multiple tissue layers and cell types. **A)** We used the µBrain preprocessed fetal gene expression dataset^97^ to parse trends in gene expression over cortical regions, cortical tissue layers, and time. We retained gene expression for 20 cortical regions and 5 tissue layers across the 16 post-conception weeks (pcw) and 21 pcw time points. We first Z-scored expression within each gene and gestational time point to represent the relative spatial localization of each gene’s expression. This allowed us to collect a spatiotemporal expression profile for each gene for subsequent clustering analysis of sulcal complexity genes. **B)** 15 of the genome-wide MAGMA significant sulcal complexity genes we identified from GWAS passed prior quality control for inclusion in the µBrain dataset, 5 of which were from the experiment-wide MAGMA significant gene set (bolded). *K*-means clustering of the spatiotemporal profile x genes matrix yielded three clusters or modules of expression of sulcal complexity genes (see also **Supplementary** Fig. 7). **C)** Sulcal complexity genes belonging to each module are shown above their module’s average spatiotemporal expression (weighted by each gene’s degree in the above clustering, **Methods**). Relative expression values are capped at Z ∈ [-1.5, 1.5] to highlight expression differences between cortical regions, tissue layers, and time points. Each module’s expression profile was correlated with the expression profiles for all 10,024 genes in the µBrain dataset, enabling ranking of all genes’ co-expression with each module for GSEA. All enrichments shown were significant following false discovery rate correction (FDR *q* < 0.05). Together, the grouping of sulci across modules suggests shape variation for both linear and complex sulci is regulated by genes expressed in germinal zones as well as the cortical plate. Source Data are provided as Source Data files.

15 of the 33 sulcal complexity genes identified by genome-wide MAGMA significance were available for annotation in the preprocessed µBrain dataset. We examined how these genes clustered in terms of their similarity in expression over fetal cortical regions, tissue layers, and time points (**Fig. 4B**), which revealed three modules of spatiotemporal co-expression. These modules were replicable when considering other candidate sulcal complexity genes identified by GWAS (**Supplementary Fig. 7**). To facilitate biological annotation of these three expression modules for sulcal complexity genes, we ranked all µBrain dataset genes by their spatiotemporal correlation with the average expression profile for each of the three corresponding sulcal complexity gene sets in **Fig. 4B** (**Methods**) – yielding three ranked gene sets amenable to gene set enrichment analysis (GSEA^92^, **Fig. 4C**, **Supplementary Data 6-7**).

Module 1 sulcal complexity genes had peak expression in the ventricular zone across at both 16 and 21 pcw, with co-expression patterns associated with proliferative cell types—cyclic (PgS/PgG2M) and intermediate progenitor cells, radial glia, and oligodendrocyte precursor cells—as well as enrichment for the nuclear cellular compartment. Module 2 genes had peak expression in the intermediate zone, most pronounced at the 21 pcw gestational stage. Module 2’s co-expression patterns were more consistent with vascular network development in the fetal brain, associated with endothelial and pericyte cells involved in the blood-brain barrier that support the development of other radially emerging proliferative cells into the intermediate zone^93,94^. Module 3 genes had peak expression in the cortical plate, especially at 21 pcw. Module 3’s co-expression patterns were enriched for maturing excitatory neuron subtypes, associated cellular compartments, and genes whose rare genetic variations were linked to developmental disorders and epilepsy^95,96^. Together, these three reproducible spatiotemporal modules (i) are enriched for distinct cellular and molecular developmental processes, (ii) span the entire radial axis of the fetal brain, and (iii) are consistent with the observed pleiotropy of distributed genetic correlations across adult cortical regions – contributing to a mechanistic model of cortical folding wherein the dynamic integration of multiple cytoarchitectural programs and biophysical processes jointly shape the complexity of any given fold.

## Discussion

We analyzed the effects of rare and common genetic variation on human sulcal complexity. By leveraging a large, multi-site structural brain MRI sample for the study of nine neurogenetic syndromes, we were able to comprehensively map syndrome-specific and shared alterations in cortical folding that may reflect shared patterns of neurodevelopmental risk. By leveraging the UK Biobank’s population-level normative adult structural brain MRI, we further identified sources of common genetic variation contributing to sulcal complexity. Because sulcal complexity reflects the end products of fetal gyrification, mapping its genetic architecture enabled us to identify candidate biological processes that shape cortical folding *in utero*. We consider each of these new insights below.

### Sulcal complexity enables novel characterization of rare genetic variation effects on cortical geometry

First, we determined cortex-wide differences in sulcal complexity across nine neurogenetic syndromes (**Fig. 1**, **Supplementary Data 1**). We show that neurogenetic syndrome impacts on sulcal complexity capture a strikingly distinct signal from those previously seen for cortical area and thickness^72–75^, because these effects: (i) are spatially uncoupled from topographies of cortical area and thickness change, (ii) do not show the reciprocal effects of genomic deletions and duplications that have been well-established for cortical area and thickness^68–70^, and (iii) are independent of brain-size alterations that co-occur in these syndromes and heavily modify regional cortical area (**Extended Data Fig. 3**, **Supplementary Fig. 2**). This distinctiveness of sulcal complexity effects in neurogenetic syndromes compared to other imaging features points to an independent set of mechanisms for their development. We speculate that one such independent mechanism may be rare variant effects on the spatiotemporal placement of sulcal roots – which could operate independently of their effects on regional cortical surface area or thickness. Sulci in the fetal brain begin as separate dimples or roots that can merge together to form continuous folds, as is the case with the two sulcal roots that together form the continuous central sulcus^22,76,98^. Consequently, the alterations of sulcal continuity detected by complexity analysis in some neurogenetic syndromes (**Fig. 1C**) may reflect rare variant effects on the timing or placement of roots for affected sulci – potentially mediated by their effects on the connectivity and cytoarchitectonic gradients thought to shape sulcal roots^4,22,99,100^.

### Convergent effects on sulcal complexity across neurogenetic syndromes

We find that there are shared, cortex-wide alterations in sulcal complexity between different neurogenetic syndromes that partly cohere with a known fetal axis of sulcal emergence (**Fig. 2**). While shared gradients of cortical area and thickness change in neurogenetic syndromes tend to smoothly follow a broad gradient between primary sensory and limbic association cortices^72,101,102^, the shared axes of atypical sulcal complexity are strikingly not smoothly graded along the cortical sheet. Rather, these axes align with the morphological ordering of folds along the linear-to-complex ranking, and therefore also with the temporal order of prenatal fold emergence from sulcal roots (**Fig. 2C-D**). This alignment suggests a parsimonious mechanism that could account for observed neurogenetic syndrome effects on the complexity of spatially remote folds. Specifically, if sulcal complexity is partly governed by the timing of sulcal root formation—creating extremes of linear or complex morphologies at earliest and latest developmental windows—then a disease-related contraction or expansion of the temporal window for sulcal root emergence would impact the differentiation of fold complexity across the cortical sheet. Fetal cohorts of individuals with rare neurogenetic syndromes, however, are presently limited to test this hypothesis^64,103,104^.

Our study design harnesses recurrent CNVs and aneuploidies as windows into rare variant effects on sulcal complexity because their recurrent nature allows estimation of an effect size map for each genetic variant. This approach provides a statistically well grounded assay for rare variant effects on cortical folding for comparison with common variant effects.

Our combined analyses of rare and common variant effects on sulcal complexity is not just informative for mechanistic understanding of cortical folding itself, but represents a rare attempt to formally and systematically compare the spatial patterning of rare vs. common variant effects on human brain organization^60,70^.

### Common genetic variation has distributed associations with sulcal complexity that broadly shape cortical anatomy and implicate diverse developmental processes

We show that the magnitude of common variant effects on sulcal complexity – as estimated through SNP-based heritability – varies substantially between folds (**Fig. 3A**). Largest heritability estimates were seen for earlier-emerging linear folds that are under tight morphological regulation^63^, also display high heritability for their sulcal depth^16,58^, and show spatial coordination with prenatally established gradients of cortical gene expression^63,105^. Conversely, later-emerging complex folds tended to show lowest heritability estimates – implying as previously hypothesized that their highly consistent tendency to show complex morphologies across individuals reflect mechanical rather than genetic constraints^63^.

There was a notable lack of strong global correspondence between inter-fold differences in common and rare variant effects on sulcal complexity (**Fig. 3A**) – although our findings do spotlight three folds that appear to be highly sensitive to both rare and common variant effects (left and right intra-parietal fissures and the left superior temporal sulcus). There is little prior work comparing the topography of rare and common variant effects on brain anatomy, so it remains unclear if the lack of concordance we observe is a distinctive property of sulcal complexity or a more general phenomenon. The high clinical penetrance of the rare variants we study here^67,106^ may be accompanied by similarly large disruptions of anatomical configuration that fall outside the range of variation reflecting common variant effects in the general population.

Common variant effects on sulcal complexity often appear to also influence the surface area (and sometimes the thickness) of immediately adjacent cortical regions – with a striking pattern where the direction of this influence “flips” across the fold (**Fig. 3B**, **Extended Data Fig. 4**, **Supplementary Figs. 5 and 6**). This observation lends support to mechanistic theories positing a linkage between the placement of cortical folds and cortical arealization^4,63,91,100^. Specifically, sharp gradients in cortical organization (consistent with the flipping of genetic correlations around folds we report here) have been proposed as spatial guides for the mechanistic buckling or folding of the cortical sheet during its early expansion in the late-second and early-third trimesters of human gestation^21,55–57^. Strikingly, we also find that genetic influences on the complexity of a given fold can be significantly associated with focal cortical anatomy variation in regions far-removed from the fold itself. This observation could indicate a secondary influence of folds on the development of remote regions (*i.e.*, shared genetics reflecting secondary impacts of the fold), or imply distributed genetic effects that simultaneously shape folding and areal expansion in remote regions of the cortical sheet. Partial support for this distributed model for cortical morphogenesis comes from our discovery and biological annotation of specific genes that influence sulcal complexity (**Fig. 3C**, **Extended Data Fig. 7**, **Supplementary Data 5**).

We specify up to 50 genes that are associated with sulcal complexity across various SNP- and gene-level tests of common variation (**Supplementary Data 5**). Many of the most robustly associated genes are known to be involved in folding-relevant aspects of fetal brain development, including several genes that have not yet been linked to cortical folding by prior work (*e.g.*, BICC1 and SLC39A10 are implicated in cell differentiation processes in embryonic development). Annotating complexity-associated genes against maps of fetal brain gene expression provides several salient insights into causal mechanisms of human cortical folding (**Fig. 4**). First – complexity associated genes are expressed along three dominant axes of fetal gene expression that together span deep germinal layers, the cortical plate and intervening intermediate zones (**Fig. 4C**) – indicating that none of these compartments is a privileged determinant of sulcal complexity. Second, complexity-associated genes did not substantially vary in their patterns of tangential expression, nor did their clustering by fetal expression group genes according to where their associated folds lay on the cortical sheet, or when their associated folds emerge in fetal development (**Fig. 4C**). This observation echoes the mixed pattern of local and remote genetic correlations observed between sulcal complexity and cortical surface area, encouraging a highly distributed mechanistic model of cortical folding. In other words, genes regulating the complexity of a given cortical fold are also often highly expressed and pleiotropically influence the morphology of remote cortical regions.

### Limitations and future directions

The present work has several limitations. First, although sulcal patterns are thought to be stable through the life span^7,8^ – and although sulcal complexity captures shape motifs that are stable across developmental and interindividual variations in size (*i.e.*, the magnitude of individual sulcal features)^63^ – the degree of stability in our continuous measure of sulcal complexity remains to be longitudinally evaluated. New and emerging resources to study cortical folding longitudinally, such as the HEALthy Brain and Child Development (HBCD) study^107^, will help evaluate the intraindividual stability of sulcal complexity over the course of postnatal development. It will further be important to replicate and explore GWAS findings in more diverse cohorts, with current samples limited to measures from individuals with predominantly European ancestry^108^. Finally, expanded fetal imaging cohorts^109^ and animal models for cortical folding, like the ferret^110^, will be necessary to test causal relationships in early development from the mechanisms we propose here.

## Conclusion

The present work provides the first genetic maps of human sulcal complexity, identifying associations with rare and common genetic variation for each of 40 sulci across the cortex. Rare and common genetic variation impacts on sulcal complexity implicate separate, new mechanisms to be considered in models of cortical folding. We provide our comprehensive, multi-scale annotations of these data as a resource to guide future studies of sulcal formation in health and its variations in clinical states, nominating specific genes and biomechanical processes for further query.

## Acknowledgments

This research was supported by the Intramural Research Program of the National Institutes of Health (NIH). The contributions of the NIH authors are considered Works of the United States Government. The findings and conclusions presented in this paper are those of the authors and do not necessarily reflect the views of the NIH or the U.S. Department of Health and Human Services. W.E.S. is a PhD student/candidate in the NIH Oxford-Cambridge Scholars Program and is also supported by the Gates-Cambridge Scholarship. W.E.S. and A.R. are supported by the Intramural Research Program of the National Institute of Mental Health (NIH annual report number ZIAMH002949). This work received computational support from the NIH HPC Biowulf cluster (http://hpc.nih.gov) and from the National Institute of Health Research (NIHR) Cambridge Biomedical Research Centre (mental health theme). All research at the Department of Psychiatry in the University of Cambridge is supported by the NIHR Cambridge Biomedical Research Centre (NIHR203312) and the NIHR Applied Research Collaboration East of England. E.T.B was supported by ImmunoMIND Medical Research Council MR/Z50354X/1 as part of the national UKRI Mental Health Platform. The views expressed are those of the author(s) and not necessarily those of the NIHR or the Department of Health and Social Care. P.E.V. is a fellow of MQ: Transforming Mental Health (MQF17_24). Funding for the 3q29 deletion syndrome project was provided by NIH R01MH118534, awarded to S.S. and J.G.M. We are thankful for the families that generously participated in the individual studies for neurogenetic syndromes and in the UK Biobank study. The UK Biobank data was obtained under application number 22875.

## Author contributions

Conceptualization and methodology: W.E.S., R.S., P.E.V., A.R., and E.T.B.; data curation: R.S., S.L., E.L, K.D., K.K., C.S., R.B., C.C., J.H., N.L., J.M., S.S., S.J. C.B.; formal analysis: W.E.S.; writing–original draft: W.E.S., P.E.V., A.R., and E.T.B; writing–review and editing: W.E.S., P.E.V., A.R., E.T.B, R.S., S.L., E.L, K.D., K.K., C.S., R.B., C.C., J.H., N.L., J.M., S.S., S.J., C.B.

## Ethics declarations – Competing interests

E.T.B. has consulted for Novartis, GlaxoSmithKline, SR One, Sosei Heptares, Boehringer Ingelheim, and Monument Therapeutics. E.T.B. is a co-founder and stockholder of Centile Bioscience Inc.

## Methods

### Imaging samples

*Neurogenetic syndrome cohorts.* We collated a primarily adolescent and young adult structural brain MRI sample of 303 individuals with neurogenetic syndromes and 312 matched neurotypical control subjects. The neurogenetic syndrome cohorts spanned 3 chromosomal aneuploidy and 6 copy number variant syndromes, all clinically ascertained. Control subjects collected as part of each neurogenetic syndrome’s studies were matched with their respective syndromic cohort on age, sex, and site data where available (**Supplementary Table 1**) (case-control differences each *p* > 0.05). Control subjects lacked the genomic alterations that were present in the neurogenetic syndrome cases, with comparable criteria for inclusion across cohorts (see **Supplementary Table 2**). All subjects’ T1-weighted MRI data were collected on 3T scanners across multiple sites across cohorts (**Supplementary Table 2**). Note that because there is high signal-to-noise ratio in identifying sulcal paths, between-site differences are thought to not substantially impact sulcal analyses^111^, especially given the high reliability of sulcal measures^58,63^ and high impacts of neurogenetic syndromes in case-control analyses^71^. Further, scan quality as indexed by Euler Number (EN)^112^ did not significantly differ by case or control status (*p*_Bonferroni_ > 0.05).

*UK Biobank cohort.* The UK Biobank is a comprehensive database and ongoing initiative to deeply phenotype biomedical traits at population-scale. We computed all sulcal measures with a similar pipeline in the UK Biobank structural brain MRI cohort (*n* = 34,725, mean age = 64 years; 46% male)^63,108,113^. In brief, the sample included adults (age range = 45–82 years) without neurological conditions^114^ scanned across three sites with the same 3T scanner model and scanning protocol. Inclusion criteria and quality control of this data are as previously described in Snyder et al.^63^ This sample enabled us to (1) benchmark the consistency of the linear-to-complex axis of cortical folding previously identified in adults with the younger cohorts presented here, (2) compute previously described gradients of sulcal morphology (*e.g.*, linear-to-complex axis, sulcal allometric scaling) to evaluate the spatial patterning of neurogenetic syndrome sulcal alterations, and (3) subsequently leverage the genotyping in this cohort to explore common variant effects on cortical folding (retained sample with genotyping: max *n* = 28,601, mean age = 64 years; 46% male).

### Measurement of sulcal phenotypes

To promote interpretability of between-syndrome relationships, we preprocessed all neurogenetic syndrome cohorts with the same pipeline starting with subject T1-weighted MRI. T1-weighted MRI were processed with FreeSurfer (v6.0.0) to delineate boundaries of gray and white matter tissue. Intensity normalization with 6 iterations (mri_nu_correct.mni –n 6) and usage of an atlas when skull stripping (–wsatlas) were included in the FreeSurfer recon-all command to reduce intensity inhomogeneities present in some cohorts of this multi-site data, which both visually and quantitatively (*i.e.*, increased EN) improved tissue segmentation. FreeSurfer segmentation was input to BrainVISA Morphologist for extraction of sulcal objects as the skeletonized paths in the space between gyri^12,13^. Sulcal regions were automatically labeled^115^ and were merged into 40 sulci identifiable in all subjects, as in Snyder et al.^63^ We expanded upon the sulcal atlas in Snyder et al.^63^ for the superior temporal sulcus region of interest to include more of the temporal parietal junction (**Extended Data Fig. 1**) as other imaging studies have implicated this region in rare neurogenetic syndromes^101^. This expanded atlas was also applied to the UK Biobank sample to compute sulcal phenotypes.

Sulcal phenotypes were derived as in our previously developed containerized pipeline^63^. In total, we measured 8 phenotypes: sulcal complexity, average depth, depth variability, longest branch, branch span, fractal dimension, sulcal surface area, and sulcal thickness for each of 40 cortical sulci per subject. We focus this study on sulcal complexity as it is hypothesized to index early sulcal development and summarizes the joint properties of sulcal average depth, depth variability, longest branch, branch span, and fractal dimension. Sulcal complexity was calculated using the methods developed in Snyder et al.^63^ First, the above five sulcal phenotypes were Z-scored across sulci within each subject. Sulcal phenotype networks (SPNs) were generated as the {40 x 40} correlation matrix between the five sulcal phenotypes. For each neurogenetic syndrome cohort, a reference SPN was computed as the element-wise mean SPN across control subjects. The first principal component (i.e., the eigen-fold index, capturing the linear-to-complex axis of sulcal shape) of the mean control SPN was correlated with individual subjects’ SPN rows in both case and control subjects, with SPN rows representing the degree to which one sulcus is morphologically similar to all other sulci in that brain. Therefore, the subject-level sulcal complexity for a given fold indexes where that sulcus’ morphology sits on the normative linear-complex axis. For sensitivity analysis, we additionally considered computing subject-level sulcal complexity for any cohort’s SPNs as the correlation between individual SPN rows and eigen-fold index computed from each of the 8 other cohorts’ mean control SPN. We also considered using eigen-fold index from the UK Biobank cohort, given the UK Biobank’s large sample enables robust estimation of the eigen-fold index or linear-to-complex axis. This allowed us to ensure that the set of control subjects in any cohort did not skew individual sulcal complexity estimates in the cohort. All sulcal complexity scores were Fisher Z-transformed to improve normality prior to estimation of case-control differences. Sulcal surface area and sulcal thickness were also computed as in Snyder et al.^63^ as these metrics are thought to biomechanically influence cortical folding^77,78^ and are most comparable to cortical area and thickness measures frequently studied in the context of neurogenetic syndromes^72,75,101^. Sulcal surface area was computed as the mean depth * length of sulcal objects^19,63^. Sulcal thickness was derived from the distance between the gray-white matter interface and the pial surface, using volumetric sulcal parcellations (subject’s sulci Voronoi diagram) as in BrainVISA Morphologist^116^.

### Quality control of sulcal morphometrics

To improve detection of case-control differences in the rare neurogenetic syndrome data, we applied rigorous quality control of sulcal data (total *n* excluded = 187, yielding final *n* = 615, see Supplementary Fig. 8). We applied four criteria for inclusion of sulcal data in downstream analyses, incorporating manual and automated evaluation of both T1-weighted MRI and derived sulcal extractions. T1-weighted MRI quality control involved standard practices of (1) manual inspection of scans for excessive head motion artifacts including ghosting, ringing, and blurring^117,118^, and (2) quantification of scan quality with a threshold of EN > −217. This threshold, representing segmentation quality which depends on scan quality, has been used previously to automatically identify scans in control and patient populations with excessive head motion or artifact^101,112^. Since artifacts can be additionally generated by poor segmentation of tissue and reconstruction of sulcal objects, we also performed (3) manual inspection of reconstructed sulci overlaid on the pial surface and (4) exclusion of subjects without all 40 sulci labeled in subject scans. Requiring all 40 sulci to be present allows for pairwise sulcal correlations in {40 x 40} SPNs and filters out subjects with poor sulcal labeling. Due to the scale of the cohort, UK Biobank sulcal data were subject to automated quality control using EN and sulcal labeling criteria as in Snyder et al.^63^

### Quantifying syndromic effects on sulcal morphology

Across each of 9 syndromes, 8 sulcal phenotypes, and 40 cortical sulci, we estimated the effect of neurogenetic syndrome diagnosis. Linear models in R (https://www.r-project.org) were used to compute the standardized effect size as the beta coefficient for diagnostic group status associated with the standardized (across individuals) sulcal phenotype (*z*_sulc_), evaluating significance with two-sided tests. Across all cohorts, we covaried for age, total tissue volume (TTV) of the cortex, and EN. Sulcal complexity is not thought to substantially vary with age^63^; however, we account for age here given sulcal complexity has not yet been explored in adolescent and young adult populations. Additionally, neurogenetic syndromes are known to have strong global effects on the brain, such as macro- and microcephaly^19,70,74^. We therefore chose to include TTV as a covariate to capture the local patterning of morphology that is often shared between syndromes^72,74,80,101^, reflecting overlapping patterns of developmental vulnerability. Standardized effect sizes were also computed without TTV as a covariate for sensitivity analysis. We used EN as a covariate as it is the standard approach to mitigate mild head-motion or scan quality effects on phenotypes^101,112,119^. Finally, where available or appropriate for each cohort, we included sex and site as covariates. The full model was specified as follows:

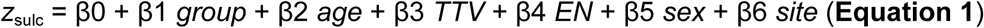

This full model was used for the 22q11.2 deletion and duplication cohorts, collected across two sites and including males and females. Site was not considered for XXY, XYY, 3q29 deletion, 11p13 deletion, and 16p11.2 deletion and duplication syndromes. Sex was not considered for XXY and XYY cohorts as 46,XY males were controls. Standardized effect sizes were derived as the beta coefficient associated with neurogenetic syndrome diagnosis from the above linear models (*i.e.*, β1), such that positive effects were associated with increases in sulcal complexity in each case group. Standardized effects were visualized on maps of cortical sulci (https://github.com/willsnyder12/sulcal_phenotype_networks). Spatial associations between effect maps and sulcal gradients used two-sided spin-based permutation with 10,000 spins to account for potential spatial auto-correlation^120^. Spins were performed using a previously developed^63^ surface-based parcellation of sulcal regions with parcel borders drawn about gyral peaks surrounding each of the 40 sulci studied here (https://github.com/willsnyder12/sulcal_phenotype_networks).

In testing for contractions of the linear-to-complex distributions of sulci in each neurogenetic syndrome cohort, sulcal complexity scores were first residualized (*sulc_resid_*) for all covariates considered (as above) other than diagnostic group. Sulci were split into linear and complex classes based on prior annotations^63^, enabling testing of opposing effects on linear and complex sulcal classes due to neurogenetic syndrome, including subject as a random intercept (*u*_subject_). This model was specified as follows:

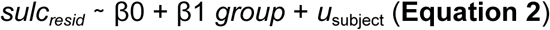

This model was fit separately for linear and complex sulcal classes, such that contraction was defined as case-control differences resulting in both increases in complexity scores for linear sulci (β1 > 0, one-sided test) and significant decreases in complexity scores for complex sulci (β1 < 0, one-sided test).

### Filtering and quality control of genetic data

In order to link inter-individual variability in sulcal complexity to inter-individual variability in the genome, additional quality control is required to mitigate spurious associations with variant-level data in the UK Biobank. We first obtained quality controlled genetic data as reported in Shafee et al.^121^ We retained subjects with non-Hispanic European ancestry to reduce bias introduced by population structure, and also excluded subjects for excessive heterozygosity, mismatches in data for genetic sex and self-reported sex, or excessive relatedness. We used imputed genetic data^122^, with variant-level filtering as previously described^121^. Altogether, 17.38 million single nucleotide polymorphisms (SNPs) were considered in subsequent analysis.

UK Biobank subjects retained from rigorous sulcal data quality control were intersected with the set of subjects retained from filtering and quality control of genetic data^121^. Sulcal complexity scores for each sulcus, computed as SPN row correlation or coherence with the linear-to-complex axis, were filtered and corrected following from standard practices in recent GWAS. For each of 40 sulci, we performed:

- Fisher Z-transformation of sulcal complexity scores to improve normality and interpretation of sulcal complexity scores at extremes
- Exclusion of outlier sulci, > mean ±5 s.d. (mean n excluded across sulci = 8.8, range = 0-66)^60,121^
- Rank-based inverse normal transformation^21,88,121,123^
- Residualisation for effects of age, age^2^, sex, age*sex, age^2^*sex, scanning site, and Euler number (EN)^60,112,117,121^

A maximum of 28,601 subjects were considered for subsequent genomic analyses. Given we found that total-tissue volume (TTV) does not substantially impact relationships between rare genetic variation and sulcal complexity, we hypothesized that it would not impact sulcal complexity associations with common genetic variation here. Therefore, we focus GWAS results on TTV-uncorrected sulcal complexity data. Providing raw, uncorrected phenotypes (such as in Warrier et al.^60^) is also important for broader utility in the field given this is the first GWAS on sulcal complexity. Further, this enables future research to have more methodological flexibility in handling global variables when integrating our GWAS results with their work (summary statistics available prior to publication). Nevertheless, for thoroughness, we computed TTV-corrected sulcal complexity, testing for congruence between TTV-corrected and uncorrected genome-wide associations in sensitivity analyses.

### Genome-wide associations

Genome-wide associations were computed with PLINK (v2.0)^124,125^. Linear models estimated additive allelic effects on sulcal complexity, covarying for the first 10 principal components of genetic ancestry^126^. Associations were computed between all 40 sulci and quality-controlled SNPs. Full dosage compensation models were used to estimate SNP associations on the X chromosome, given most indices for cortical morphometry are best explained by a full dosage compensation model^127^. To adjust significance for correlated tests, we calculated the number of effective tests from the full 40 sulci using the approach from Nyholt^128^, resulting in 34 effective tests. In subsequent analyses, we considered SNP associations at a conservative experiment-wide threshold (*p* < 5 ×10^−8^ / 34, or, *p* < 1.47 ×10^−9^), in addition to the standard threshold of genome-wide significance (p < 5 ×10^−8^). Lead SNPs for independent loci were identified per sulcus using FUMA with default linkage disequilibrium (LD)^129^ of *r*^2^ < 0.1. SNP-based heritability (*h*^2^) for each sulcus was computed with Linkage Disequilibrium Score Regression (LDSC)^81^ with one-sided significance evaluated (*h*^2^>0).

To map from SNP to gene-level associations, we considered two standard approaches. First, we applied MAGMA (Multi-marker Analysis of GenoMic Annotation, v1.1) with a 10kb symmetric window to aggregate SNP-level effects to proximal protein coding genes^82^. MAGMA has been shown to reduce false positives, account for LD structure, and has direct interpretation of genomic variation impacting the sequence of expressed transcripts or structure of proteins or that of their nearby regulatory regions^82^. Gene-level significance involved correction for effective tests, for which we considered genes at experiment (*p* < 0.05 / 19,299 genes / 34 effective tests, or, *p* < 7.62 ×10^−8^) and genome-wide thresholds (*p* < 2.59 ×10^−6^). As a complementary approach to positional mapping with MAGMA, we additionally identified genes proximal (within a 10kb symmetric window) to any individually significant SNP, supporting identification of genes where more focal SNP effects may be relevant. Genes proximal to SNPs with associations reaching experiment (*p* < 1.47 ×10^−9^) and genome-wide significance (*p* < 5 ×10^−8^) were considered. As a limitation to this present study, current annotations of SNP-effects to genes leveraging chromatin interaction profiles^130^ or expression quantitative trait loci^131^ do not have the necessary fetal brain gene expression data for the specific developmental windows or tissues associated with cortical folding and were therefore not considered. Significant genes identified using either the MAGMA or SNP-based approaches, at either experiment or genome-wide significance, were all considered for gene-ontology enrichment using WebGestalt^132^. Gene lists of either the full genome or only protein-coding genes in the brain (from Wagstyl et al.^105^) were used as background genes in over-representation analysis. Additionally, we compiled summaries for all significant genes using biomaRt^133^ and National Center for Biotechnology Information (NCBI) Entrez^134^ resources. For select loci lying in intergenic regions, we visualized proximal genomic regions alongside SNP effects, LD structure, and recombination rates using locuszoomr^135^.

### Multi-scale annotations of sulcal complexity genetics

Genomic associations were comprehensively annotated against a variety of imaging, genomic, and transcriptomic resources.

### Axes of normative and syndromic sulcal morphology

To evaluate whether the magnitude of common variant influences on sulcal complexity (estimated as *h*^2^ from GWAS) was associated with a sulcus’ position on the linear-complex axis or its sensitivity to rare variant effect, we determined the cross-sulcal spatial correlations between these sulcal maps. Spatial correlations between sulcal maps were corrected for spatial auto-correlation with spin-based permutation with 10,000 spins^120^.

To assess genetic architecture shared between sulcal complexity and other phenotypes, we estimated genetic correlation (r_g_) with cortical morphometry, and multiple psychiatric, behavioral and development traits. Cortical surface area and thickness were selected as extensively studied cortical morphometry measures hypothesized to be mechanistically linked to cortical folding. Psychiatric diagnoses in addition to behavior and development phenotypes were selected for their hypothesized associations with early life development or broad influence on cortical structure. Genetic correlations were computed using LDSC with reference European-ancestry LD scores from the 1000 Genomes Project^81^. Summary statistics were cleaned (munge_sumstats.py) and filtered for HapMap3 SNPs^81,136^. The following prior GWAS summary statistics were used for genetic correlations with all 40 sulci:

- Cortical morphology (Shafee et al.^121^)

– surface area, measured on 360 regions in the Glasser^3^ parcellation

– thickness, measured on 360 regions in the Glasser^3^ parcellation

- Neuropsychiatric disorders

– ADHD (Demontis et al.^137^)

– Alzheimer’s disease (ALZ) (Jansen et al.^138^)

– Autism spectrum disorder (ASD) (Autism Spectrum Disorders Working Group of The Psychiatric Genomics Consortium^139^)

– Bipolar disorder (BP), type I and type II cases, including UK Biobank data (Mullins et al.^140^)

– Major depressive disorder (MDD), excluding 23andMe data (Wray et al.^141^)

– Schizophrenia (SCZ), primary autosome data (Trubetskoy et al.^142^)

- Behavior and development

– Educational attainment (Okbay et al.^143^)

– Intelligence (Savage et al.^144^)

– Age at onset of walking (AoW) (Gui et al.^145^)

– Head circumference at birth (Vogelezang et al.^146^)

– Birth weight, fetal effect, European-ancestry sample (EGG Consortium et al.^147^)

– Gestational duration, fetal effect (Liu et al.^148^)

### Lead SNP associations

To determine if specific common variants influencing sulcal complexity are also associated with other aspects of brain anatomy, we mapped lead sulcal complexity SNPs to previous GWAS on brain morphometry measured at sulcal resolution. Specifically, the UK Biobank multimodal brain imaging GWAS (“BIG40”)^88^ summary statistics include morphometry from the Destrieux atlas^9^, with coverage over sulcal and gyral regions comparable to the sulcal atlas in our present analysis. Lead SNPs identified from sulcal complexity GWAS were visualized for their significance for local CT and SA GWAS (**Fig. 3B**,

**Extended Data Fig. 4, Supplementary Figs. 5-6**) to gauge spatial overlap between sulcal complexity GWAS and GWAS of these features mechanistically implicated in cortical folding.

### Fetal brain transcriptomics

Finally, we explored the fetal brain expression of genes from sulcal complexity GWAS in the µBrain dataset^97^. We used preprocessed data generated from laser microdissection and RNA microarray^149^ of 4 brains from 16 to 21 post-conception weeks (pcw), *i.e.*, early to mid-gestation. This dataset is well-suited for probing mechanisms of cortical folding as it has comprehensive sampling across cortical regions and along tissue layers in these time points immediately preceding the onset of cortical folding. We subsetted data to 20 cortical regions where sulcation occurs, ordered anterior to posterior as follows: orbitofrontal, subcallosal cingulate, frontal pole, rostral cingulate, ventrolateral prefrontal cortex, dorsolateral prefrontal cortex, auditory, ventrolateral temporal, medial temporal-occipital, entorhinal, mid-cingulate, motor, sensory, caudal cingulate, dorsolateral temporal, lateral temporal-occipital, ventral parietal, dorsal parietal, primary visual, and extrastriate. Note that present fetal cortical gene expression datasets are limited by the sampling schema in fetal transcriptomics, where cortical regions were sampled without discernment over prospective sulcal or gyral territory^97,149,150^. However, most variability in gene expression appears to occur radially across cortical layers rather than tangentially across cortical regions^97^ (**Fig. 4C**), with such variance captured well by this present dataset. We retained all cortical layer data available for the cortical plate, subplate, intermediate zone, subventricular zone, and ventricular zone. Data were averaged across specimens for the ∼16 pcw time point (specimen H376.IIIA.02, H376.IIIB.02) and the ∼21 pcw time point (H376.IV.02, <u>H376.IV</u>.03).

Genome-wide significant genes determined from MAGMA were explored in the µBrain dataset, where quality-controlled data was available. For each gene associated with sulcal complexity, we Z-scored expression within each time point, allowing characterization of the relative or localized expression for each gene and its change over time points. Next, we clustered a {genes×(regions×layers×time)} matrix to identify spatiotemporal modes of gene expression during early to mid-gestation. K-means clustering was performed for *k* ∈ [2…*n*_genes_/2 ]. To provide biologically interpretable clusters that avoid over- or underfitting, *k* was chosen using the elbow criterion applied to the ratio of within-versus between-cluster sum of squared distances per value of *k*. Prototype spatiotemporal expression profiles for each cluster, or module, was estimated as the degree-weighted average expression across module genes, using gene-pairwise spatiotemporal co-expression to calculate degree. For sensitivity analysis, we recomputed and compared gene modules using all candidate sulcal complexity genes from genome-wide significant MAGMA genes and genes proximal to genome-wide significant SNPs. The spatiotemporal pattern of prototype module expression was correlated with all fetal brain expressed genes, creating a ranking of all genes by their similarity to each module’s spatiotemporal expression pattern. These gene rankings were subjected to gene-set enrichment analysis (GSEA)^92^ on fetal cell types^94^, cellular compartments^151^, and disease^95,96,152–155^ gene ontologies as compiled in Wagstyl et al.^105^

## Data availability

All genetic maps of rare neurogenetic syndrome or common variant effects on sulcal complexity produced in this study are provided in **Supplementary Data** and **Source Data** files. Additionally, genetic correlations with cortical morphometrics, gene summaries of GWAS significant genes, fetal brain transcriptomic module data and related gene set enrichment analysis are all provided in **Supplementary Data** and **Source Data** files. Sulcal complexity GWAS summary statistics will be deposited with accession codes made available prior to publication. Raw neuroimaging data for neurogenetic syndromes are available through request and data access agreement from the principal investigators of the projects from the studies they are derived from. UK Biobank imputed genotype data and imaging data can be accessed through application to the UK Biobank (https://www.ukbiobank.ac.uk/use-our-data/apply-for-access/). Sulcal gradients described in Snyder et al.^63^ were reproduced with the updated sulcal atlas used in the present study, and are also provided in **Source Data** files. Sulcal emergence data was obtained from Snyder et al.^63^ Associations with GWAS on area and thickness phenotypes were derived from Shafee et al.^121^ and from Smith et al.^88^ (https://open.oxcin.ox.ac.uk/ukbiobank/big40/). Fetal brain transcriptomics were obtained from Ball et al.^97^ (https://zenodo.org/records/10622337).

## Extended Data for

**Extended Data Figure 1.**
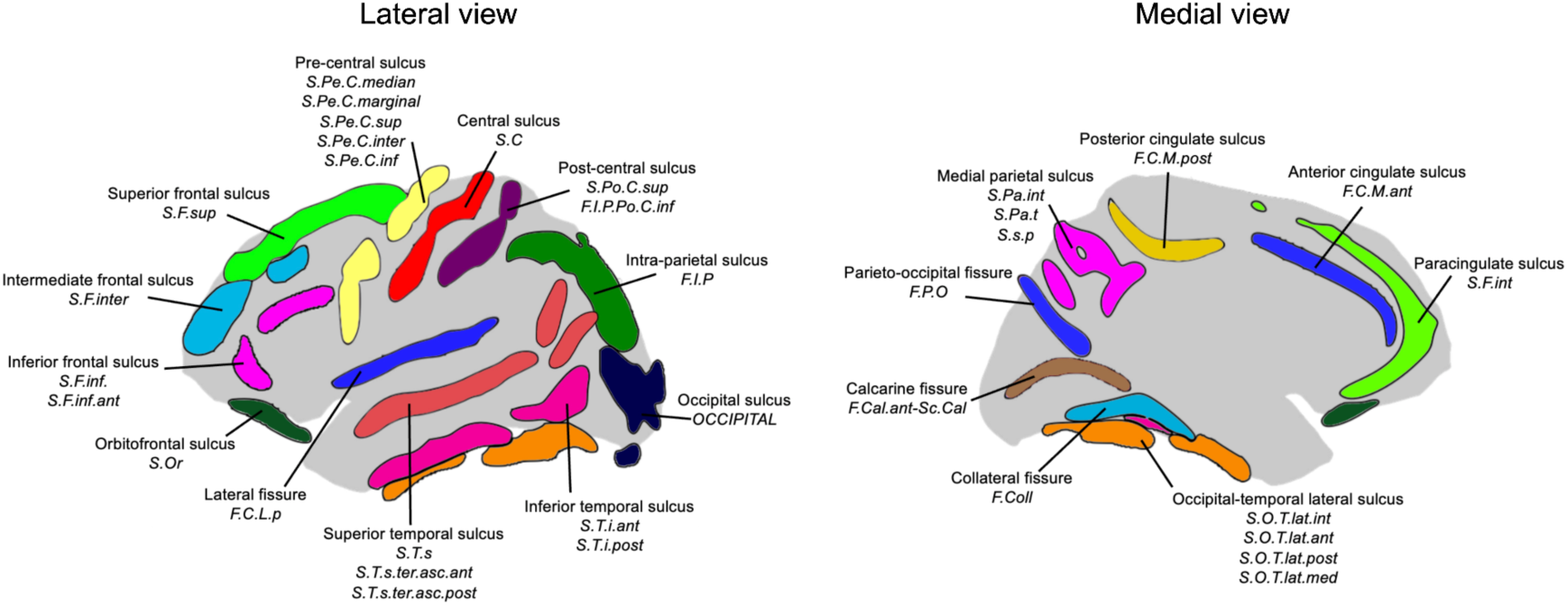
**Sulcal atlas used for labeling of sulcal regions in neurogenetic syndrome cohorts and the UK Biobank cohort**. The atlas of 40 cortical sulci spanning the cortex that was used in Snyder et al.^63^ was expanded upon to include regions in the temporoparietal junction (*S.T.s.ter.asc.ant, S.T.s.ter.asc.post*) as part of the superior temporal sulcus label.

**Extended Data Figure 2.**
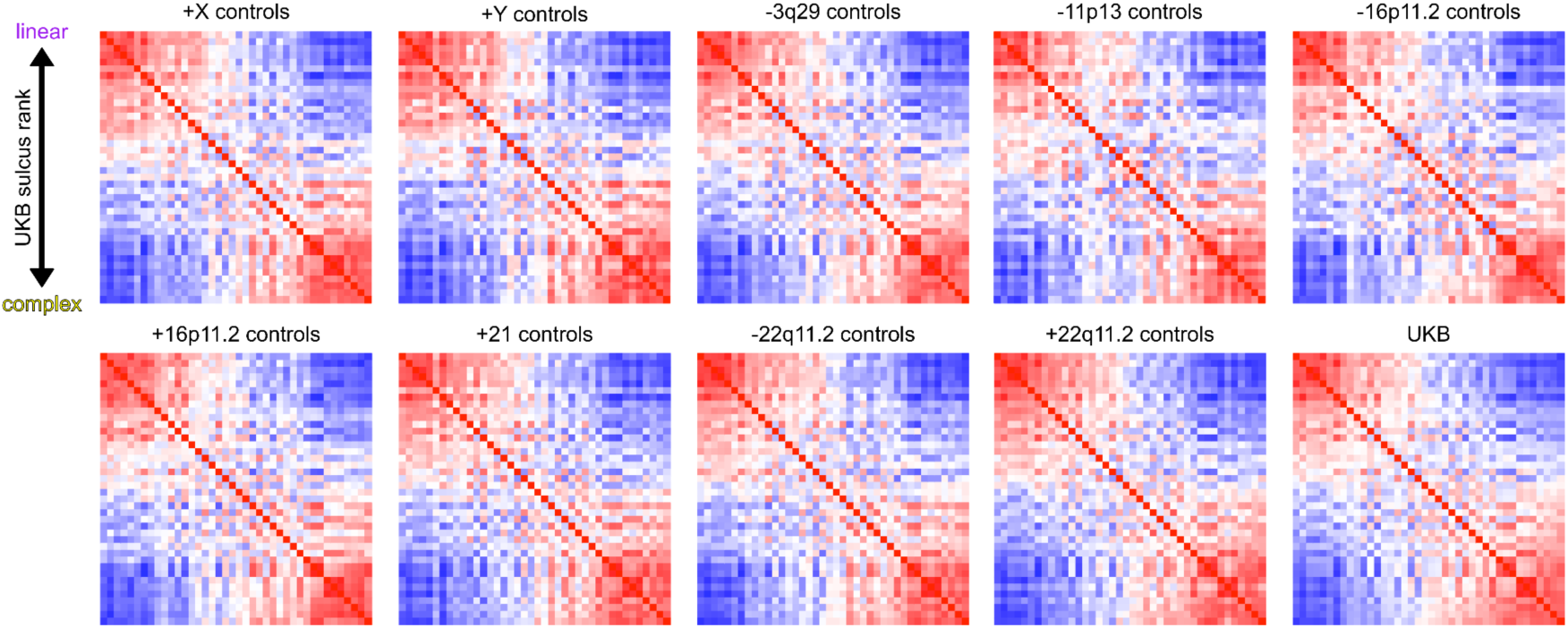
Overall sulcal architecture is conserved across neurotypical populations of varying age ranges. The {40 sulci x 40 sulci} mean SPN matrix is shown for each set of controls matched to respective neurogenetic syndrome cohorts. Sulci were arranged the same in each matrix by the rank of sulcal complexity in the UK Biobank mean SPN, given by its first principal component. The bimodal structure of sulcal morphology across cortical regions is consistent across all cohorts in terms of the edge-level, bipartite cluster-level, and row (fold)-level structure of mean SPNs. At the edge-level, mean control SPNs were highly similar between control groups matched to neurogenetic syndrome groups (mean edge-wise correlation *r* = 0.93; cohorts’ mean age range: 13-37 years) and were highly similar to the UK Biobank adults (mean *r* = 0.96, UK Biobank age range: 45–82 years), indicating that the bimodal axis of sulcal morphology (*i.e.*, consistent shape ranking along a linear-to-complex axis of shape variation) is strongly conserved across the life span. At the cluster-level, there is consistent bipartite clustering of mean SPNs across all cohorts (90% agreement in fold-level clustering into linear or complex folds as in the UK Biobank mean SPN, with 70% of sulci having 100% consistent clustering across all control cohorts). At the row (fold)-level, the relative ranking of folds along the linear-to-complex axis – derived as the first principal component of the UK Biobank mean SPN – was strongly recapitulated in each set of controls matched to respective neurogenetic syndrome cohorts (mean Spearman’s ⍴ = 0.98). Together, we demonstrate that the previously described bimodal structure of SPNs in adulthood is faithfully mirrored in neurotypical youths, providing a basis for reliable computation of sulcal complexity in each neurogenetic syndrome cohort (see also **Supplementary** Fig. 1). Source Data are provided as Source Data files.

**Extended Data Figure 3.**
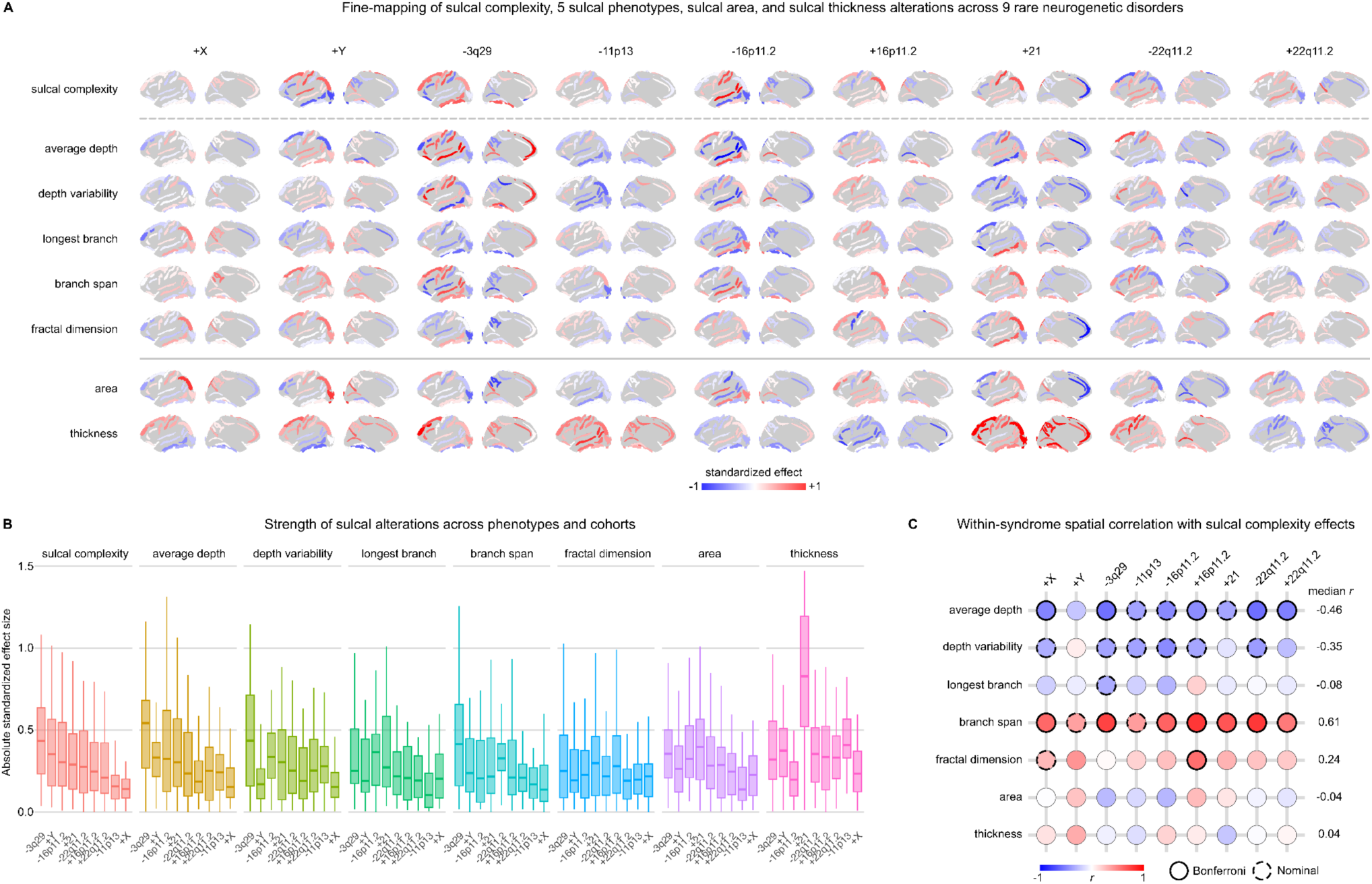
Comparing changes in sulcal complexity to alterations in constituent sulcal phenotypes. **A)** Standardized effects of 9 neurogenetic syndromes were computed for 7 sulcal phenotypes and 40 sulci, unthresholded for significance to highlight spatial patterns between syndromes and sulci. Sulcal maps are bilaterally averaged for visualization. Along columns, inspection of sulci with strong sulcal complexity effects reveals comparably strong relative effects on other sulcal phenotypes. Positive effects (red) indicate increased values in neurogenetic syndromes compared to controls, and negative effects (blue) indicate decreased values for these sulcal phenotypes. Strong increases in sulcal complexity were often seen coinciding with decreases in average depth and increases in branch span for the same sulcus, with the opposite trend observed with strong decreases in sulcal complexity. See **Supplementary Data 1** for all effect sizes and associated *p*-values. **B)** Distributions of absolute effect sizes across cortical sulci are plotted for each phenotype and for each syndrome. Box plots are ordered for each phenotype by the rank of median strength of sulcal complexity effects. The relative ranking of neurogenetic syndromes by average absolute effect size is similar for sulcal complexity and other sulcal phenotypes, except for sulcal thickness. **C)** Spatial correlations were determined between each phenotype’s standardized effects and sulcal complexity standardized effects for each syndrome. Significance is corrected for tests across syndrome and adjusted for effects of spatial auto-correlation (Bonferroni: *p*_spin_ < 0.05/9, nominal: *p*_spin_ < 0.05). Average depth and branch span are most linked to the spatial patterning of sulcal complexity. Sulcal surface area and sulcal thickness are inconsistent in the sign of their relationships with sulcal complexity across syndromes and did not show any significant shared spatial patterning. Source Data are provided as Source Data files.

**Extended Data Figure 4.**
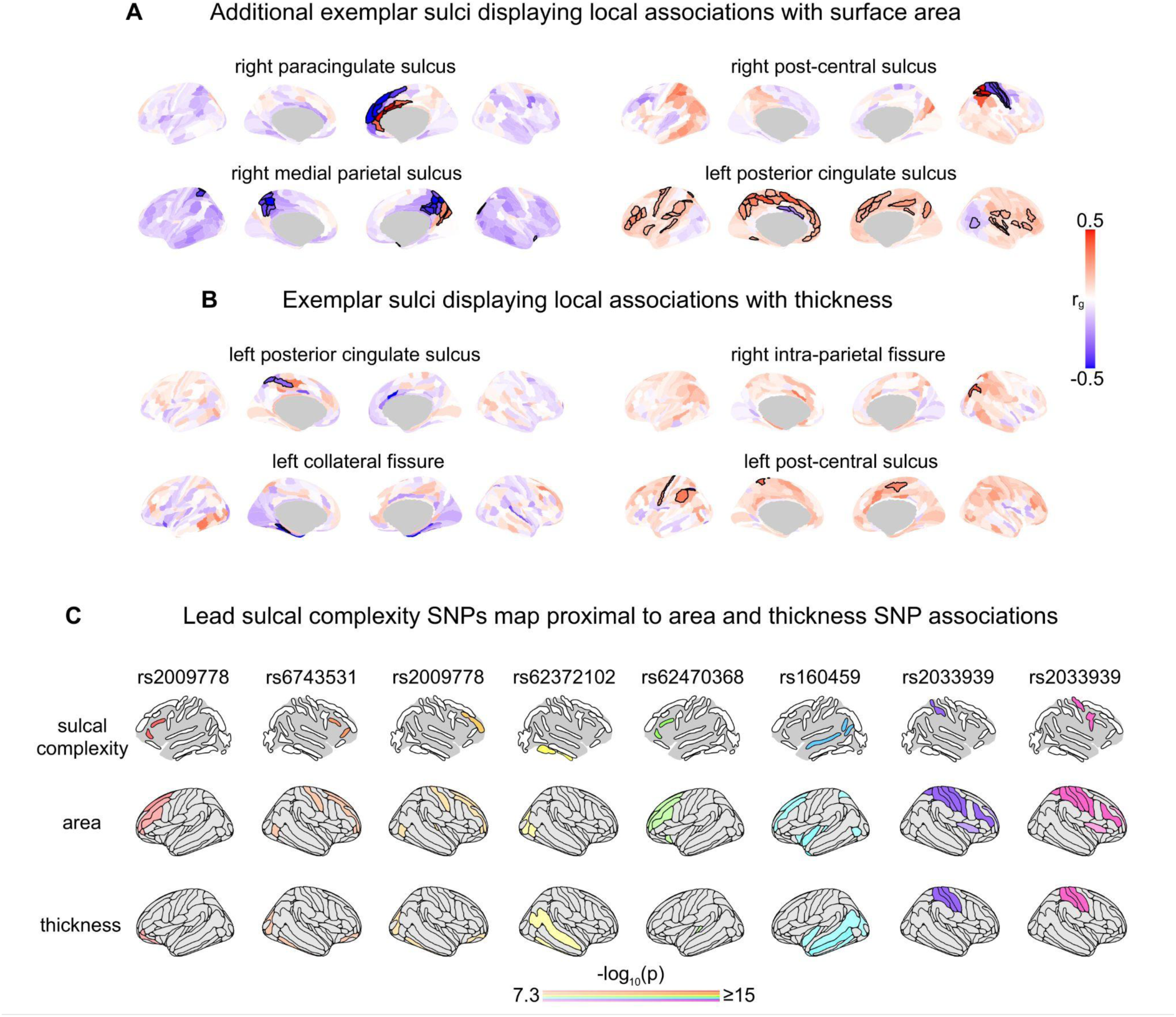
Sulcal complexity genetic correlations with surface area and thickness demonstrate local and global spatial patterns of shared genetic architecture. **A)** Expanding on Fig. 3B, we display additional exemplars of genetic correlations between sulcal complexity and cortical surface area throughout the brain. Again, patterns of spatially local and remote genetic correlations with each fold are observed. Spatially local genetic correlations flip sign proximal to the location of each fold for which significant genetic correlations are observed, indicating a genetic gradient for cortical arealization is linked to variation in fold complexity. Significant positive and negative associations are bolded and corrected across cortical parcels (FDR *q* < 0.05). All genetic correlations between cortical surface area and each sulcus’ sulcal complexity are shown in **Supplementary** Fig. 5, **Supplementary Data 4**. **B)** Similar to the above associations, spatially local genetic correlations can also be seen for cortical thickness – wherein the common genetic variation predicting increased sulcal complexity can either predict increased or decreased cortical thickness, depending on the sulcus. Significant positive and negative associations are bolded and corrected across cortical parcels (FDR *q* < 0.05). Thickness associations appear to be more spatially local to the sulcus than area genetic correlations, in that significant regions for thickness genetic correlations tend to only overlap with the sulcus of interest. All genetic correlations between cortical thickness and each sulcus’ sulcal complexity are shown in **Supplementary** Fig. 6. **C)** Lead SNPs from independent significant loci identified in sulcal complexity GWAS (top row) were investigated for SNP effects on local surface area and thickness in prior GWAS (Smith et al.^88^). The lead SNPs’ effects are plotted for its associations in each cortical region’s separate GWAS, with regions not reaching genome-wide significance colored gray (p < 5 ×10^−8^) and color saturation adjusted for values capped at −log(*p*) = 15 for visualization. Lead sulcal complexity SNPs, associated with specific sulci, are also strongly associated with surface area and thickness in cortical regions proximal to these sulci. Therefore, not only is the distributed common genetic variation shared between local sulcal complexity and area or thickness, but also specific loci with the strongest genetic associations are shared between local sulcal complexity and area or thickness. Source Data are provided as Source Data files.

**Extended Data Figure 5.**
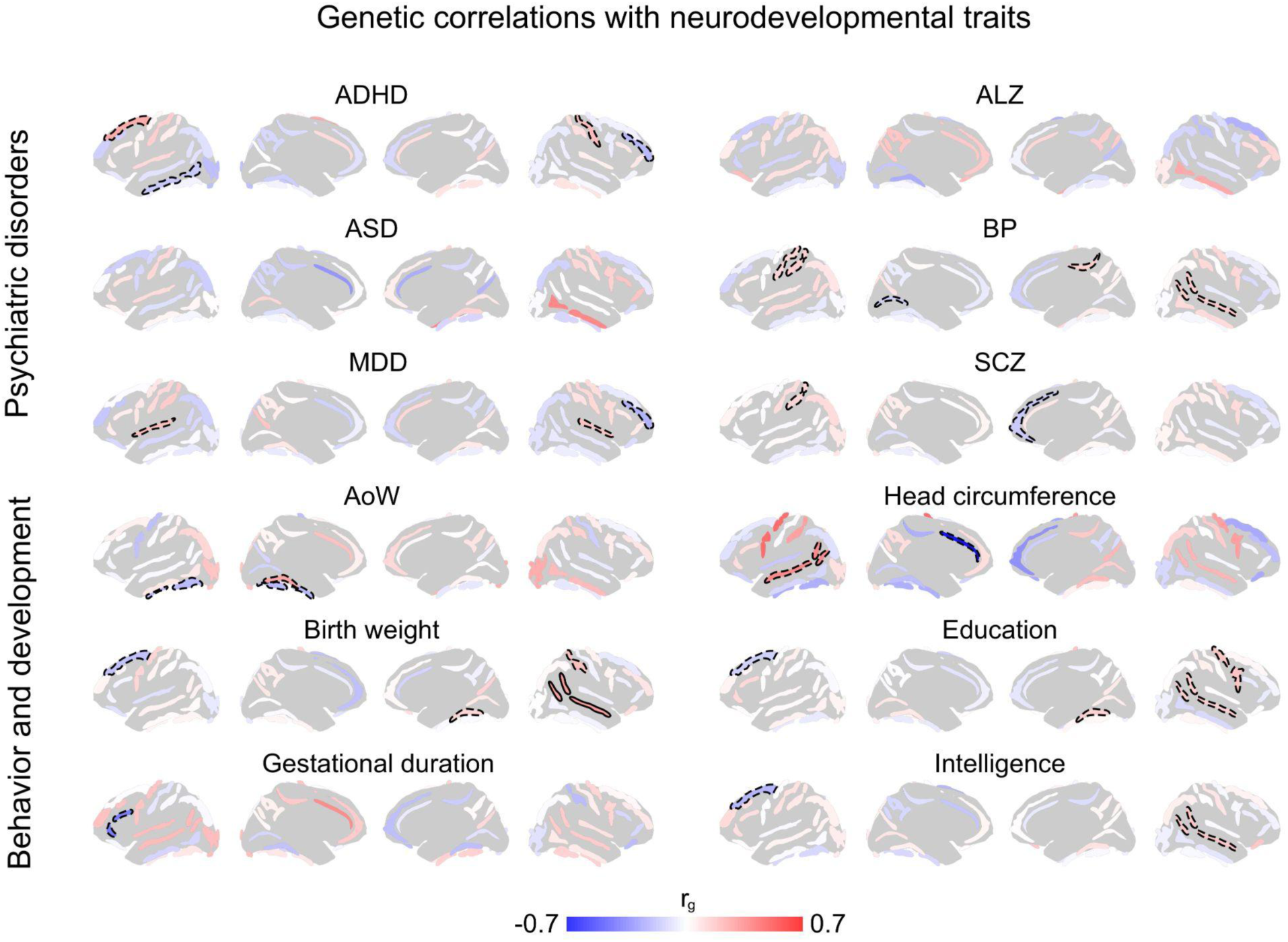
Limited evidence for shared genetic architecture between sulcal complexity and neurodevelopmental traits. Six psychiatric and disease traits were tested for genetic correlations with sulcal complexity across 40 sulci. Only nominally significant (dashed outline, p < 0.05) genetic correlations were observed, with different sets of sulci across disorders/diseases. Additionally, six behavioral and developmental traits were tested for genetic correlations. Associations were again limited across sulci. Only one significant association was found (bold outline, *p* < 0.05 / 34 effective sulci), demonstrating positive genetic correlation between birth weight and right superior temporal sulcus sulcal complexity. Source Data are provided as Source Data files.

**Extended Data Figure 6.**
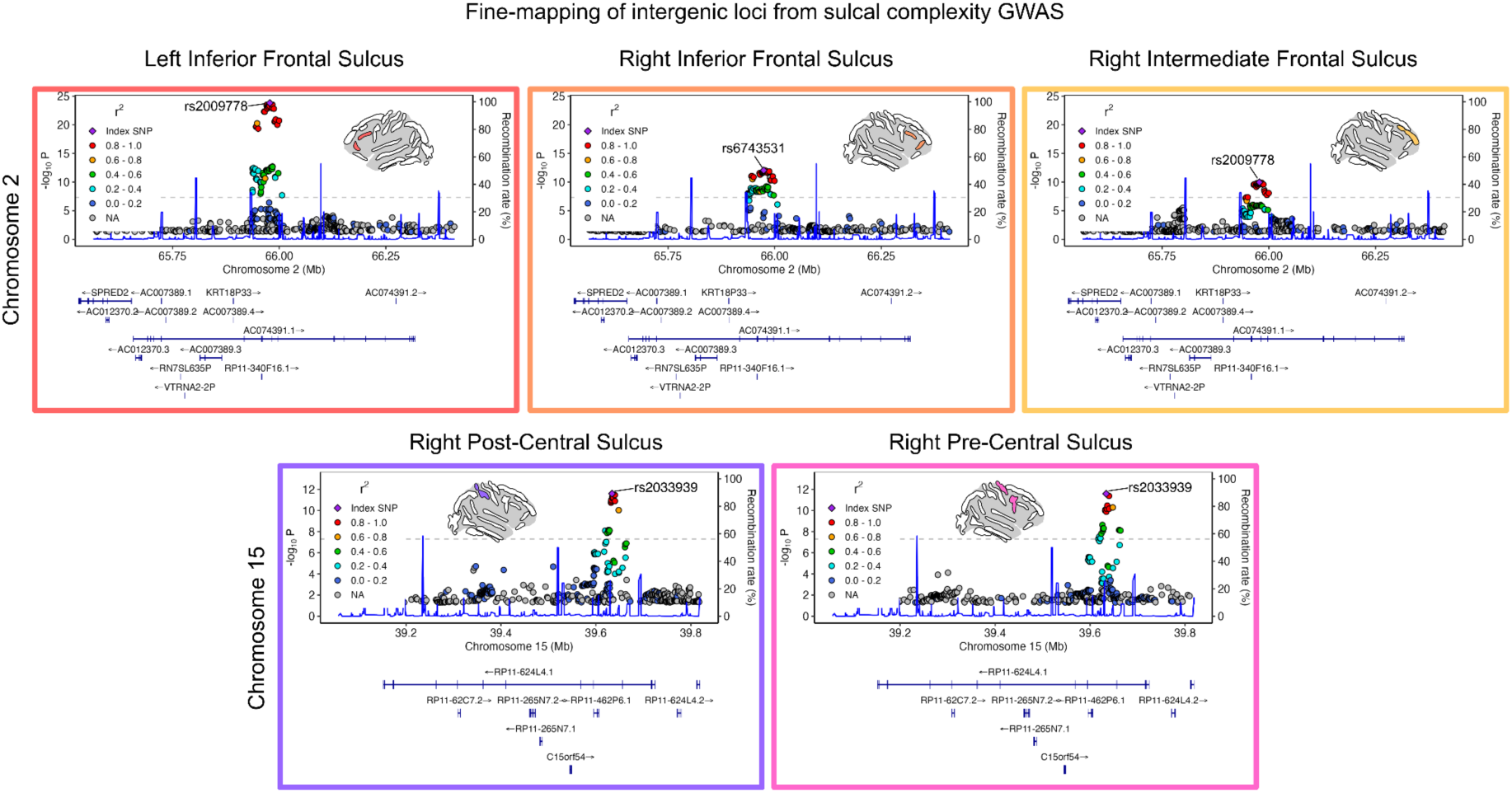
Fine-mapping of intergenic loci from sulcal complexity genome-wide associations. Gene tracking is shown for loci identified from sulcal complexity GWAS that did not map to protein-coding genes. For all locuszoomr plots, genomic regions, recombination rate, and linkage disequilibrium (LD) are shown with SNP associations with each sulcus. (Top) The locus on chromosome 2 overlapped for the left inferior frontal sulcus, right inferior frontal sulcus, and right intermediate frontal sulcus, which were all contained within the long intergenic non-coding RNA (LINC) region AC074391.1 (LINC02934). (Bottom) The locus on chromosome 15 overlapped for the right post-central sulcus and right pre-central sulcus, which were all contained within the LINC region RP11-624L4.1 (LINC02915).

**Extended Data Figure 7.**
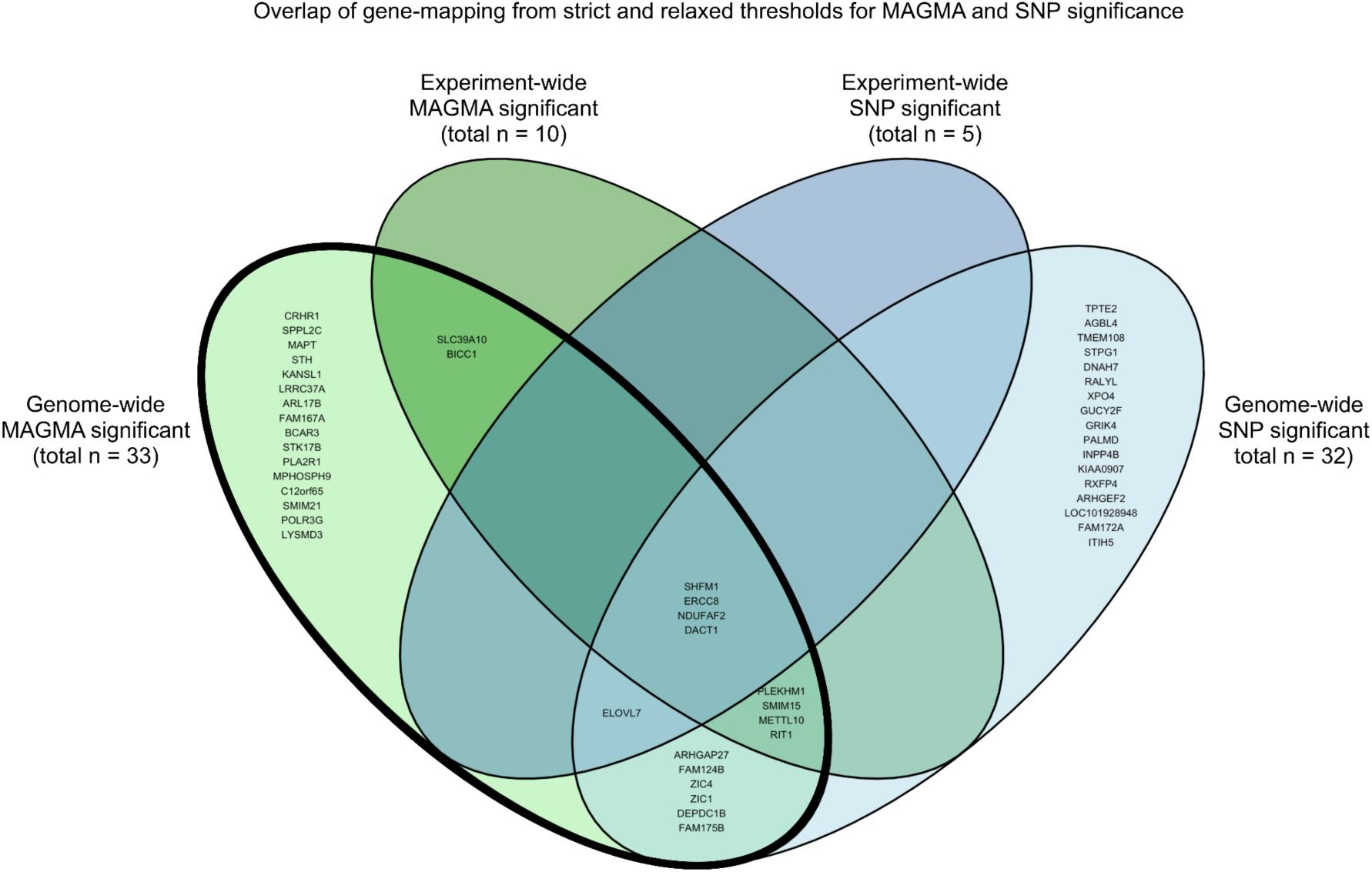
Candidate sulcal complexity gene sets converge with different approaches. Two approaches at two thresholds were used to determine candidate sulcal complexity genes from GWAS. In main results, we focus on the genes identified by MAGMA, as MAGMA has been well-validated for use with GWAS and captures summative effects of SNPs proximal to genes. Experiment-wide significance corrected for the number of protein-coding genes and effective number of sulci tested (*p* < 7.62 ×10^−8^) and genome-wide significance corrected for the number of protein coding genes (*p* < 2.59 ×10^−6^). In subsequent analyses in fetal transcriptomic data, we focus on the genome-wide MAGMA significant gene set (bold outline) to retain an expanded set of candidate genes. Alternatively, we also identified genes that were proximal to individually significant SNPs. Genes within 10kb of experiment-wide significant SNPs (correcting for genome-wide SNP threshold and number of effective sulci, *p* < 1.47 ×10^−9^) and genome-wide significant SNPs (*p* < 5 ×10^−8^) were considered. Overall, these methods provide complementary identification of 50 candidate sulcal complexity genes.

