## Supplementary Figures 1-8. Supplementary Tables 1-2. for "Genetic insights on the mechanisms of human cortical folding": Supplementary_Information_Snyder_et_al_2026.pdf

1. Section on Developmental Neurogenomics, Human Genetics Branch, National Institute of Mental Health Intramural Research Program, Bethesda, MD, United States of America
2. Department of Psychiatry, University of Cambridge, Cambridge, UK
3. Lifespan Brain Institute, Children's Hospital of Philadelphia and Penn Medicine, Philadelphia, Pennsylvania, United States of America
4. Marcus Autism Center, Children's Healthcare of Atlanta, Atlanta, Georgia, United States of America, Emory University School of Medicine, Department of Pediatrics, Atlanta, Georgia, United States of America
5. Emory University School of Medicine, Department of Pediatrics, Atlanta, Georgia, United States of America
6. Tri-Institutional Center for Translational Research in Neuroimaging and Data Science (TReNDS), Georgia State University, Georgia Institute of Technology and Emory University, Atlanta, Georgia, United States of America
7. Centre de recherche CHU Sainte-Justine and University of Montreal, Canada
8. Department of Psychiatry and Biobehavioral Sciences, Semel Institute for Neuroscience and Human Behavior, University of California, Los Angeles, Los Angeles, California, United States of America
9. Imaging Genetics Center, Mark and Mary Stevens Neuroimaging and Informatics Institute, Keck School of Medicine of the University of Southern California, Marina del Rey, Los Angeles, California, United States of America
10. Division of Pediatric Endocrinology, Nationwide Children's Hospital, Ohio State University, Columbus, Ohio, United States of America
11. Department of Psychological and Brain Sciences, Drexel University, Philadelphia, Pennsylvania, United States of America
12. Department of Psychiatry, Robert Wood Johnson School of Medicine, Rutgers University, United States of America
13. Center for Advanced Biotechnology and Medicine, Rutgers University, United States of America
14. Department of Pediatrics, University of Montreal, Montreal, QC, Canada
15. Department of Psychology, University of California, Los Angeles, CA, United States of America
16. School of Academic Psychiatry, Institute of Psychiatry, Psychology & Neuroscience, King's College London, London, UK
* These authors contributed equally

### Table of Contents

**Supplementary Figure 1.** Individual estimates of sulcal complexity are not biased by the control sample matched to each neurogenetic syndrome cohort.
**Supplementary Figure 2.** TTV-uncorrected sulcal alterations are spatially similar to TTV-corrected sulcal alterations.
**Supplementary Figure 3.** The shared axis of ∆complexity effects is replicable with TTV-uncorrected ∆complexity estimates.
**Supplementary Figure 4.** TTV regression effect on sulcal complexity heritability.
**Supplementary Figure 5.** Genetic correlations between sulcal complexity and regional surface area.
**Supplementary Figure 6.** Genetic correlations between sulcal complexity and regional cortical thickness.
**Supplementary Figure 7.** *K*-means clustering and reproducibility of sulcal complexity gene modules in fetal spatiotemporal gene expression data.
**Supplementary Figure 8.** Quality control (QC) of imaging data from neurogenetic syndrome cohorts.
**Supplementary Table 1.** Demographics of each neurogenetic syndrome cohort.
**Supplementary Table 2.** Additional cohort information on ascertainment, inclusion criteria, and scanning protocols.
**Description of Additional Supplementary Data Files.** Descriptions of Supplementary Data 1-8.

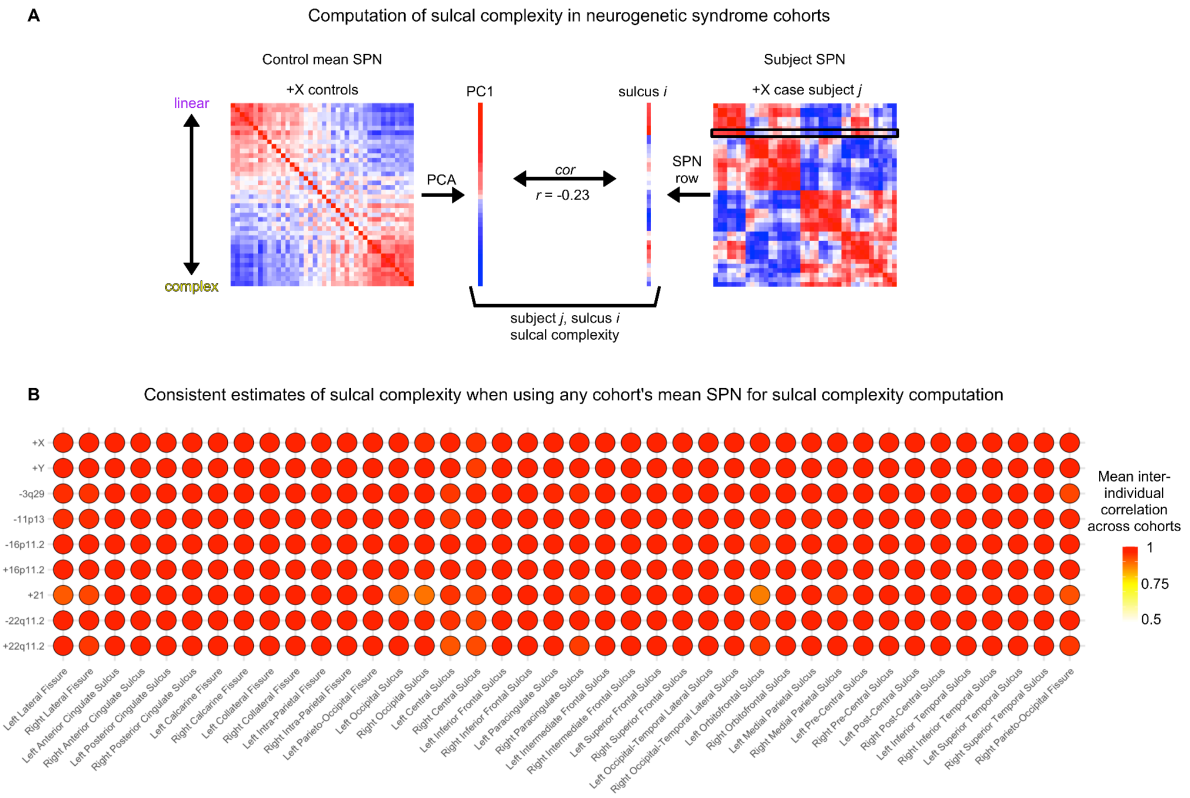


Supplementary Figure 1. **Individual estimates of sulcal complexity are not biased by the control sample matched to each neurogenetic syndrome cohort.**

**A)** Sulcal complexity is derived from sulcal phenotype networks (SPNs) as the coherence between an SPN row and the first principal component (PC1, from principal component analysis, PCA) of the mean SPN of each control group matched to neurogenetic syndrome cohorts. Displayed is an example of the computation of one sulcus' sulcal complexity for an individual with XXY diagnosis using the mean SPN from controls matched to individuals with XXY diagnosis. To ensure sulcal complexity had comparable interpretation across cohorts (*i.e.*, no bias based on sampled control group), we recomputed sulcal complexity in each cohort with the mean control SPN from control groups in each other of the 9 syndromic cohorts and when using the UK Biobank mean SPN. **B)** Displayed are the average inter-individual correlations between (1) sulcal complexity derived using a cohort's matched controls and (2) sulcal complexity derived using any of the control cohorts' or UK Biobank's mean SPN. Strong positive correlations for all syndrome cohorts and for each sulcus (across syndromic cohorts' control mean $r$ = 0.98, using UK Biobank as reference mean SPN $r$ = 0.97) indicate high replicability of sulcal complexity, enabling reliable interpretation of cross-syndrome sulcal complexity.

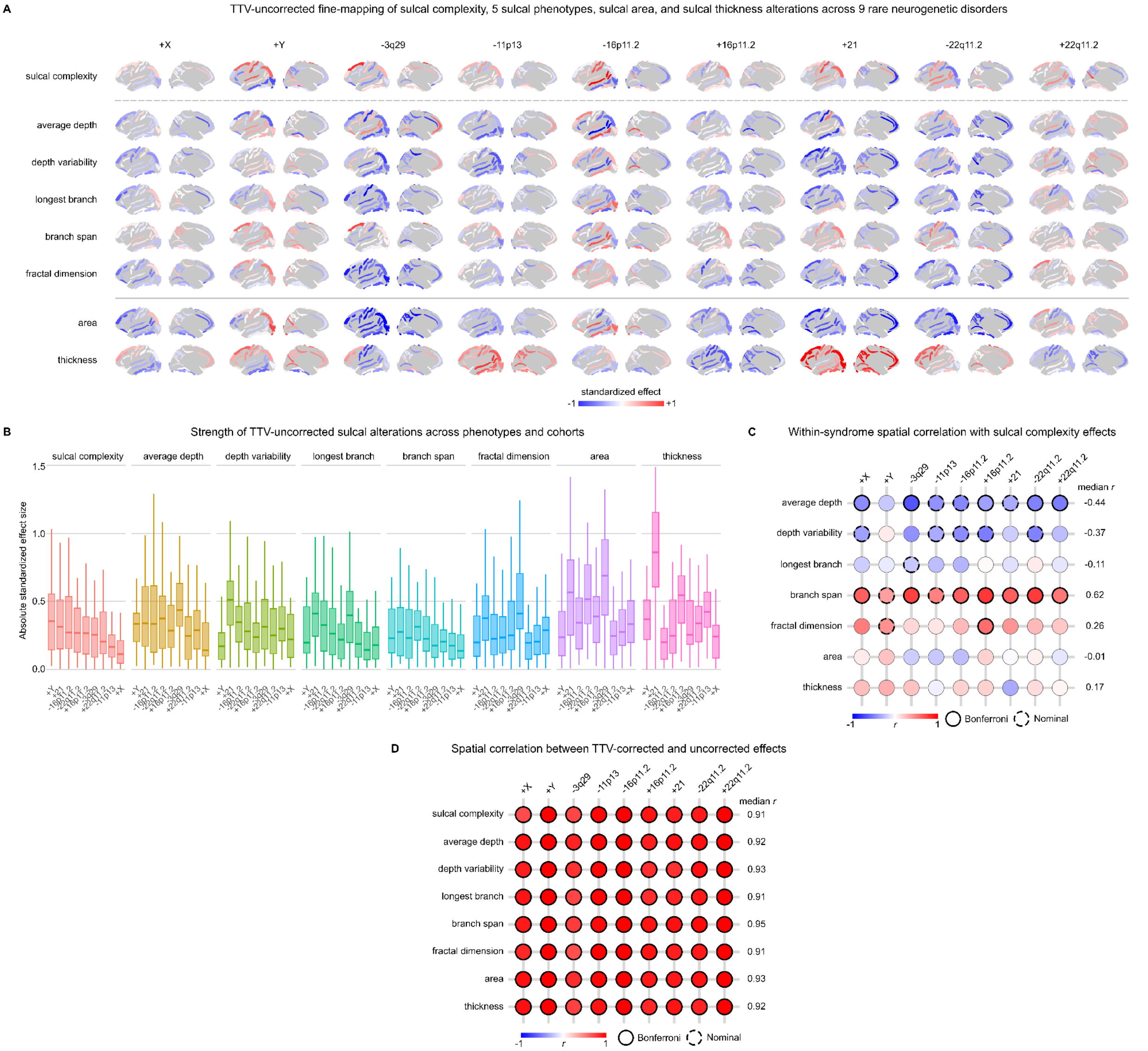


Supplementary Figure 2. **TTV-uncorrected sulcal alterations are spatially similar to TTV-corrected sulcal alterations.**

**A)** Standardized effects of 9 neurogenetic syndromes were recomputed for 7 sulcal phenotypes and 40 sulci without covarying for total tissue volume (TTV). Sulcal maps were bilaterally averaged for visualization. Qualitatively, the sign of TTV-uncorrected sulcal complexity effects is almost identical to that of TTV-corrected sulcal complexity effects (see **Extended Data Fig. 2** for TTV-corrected effects). However, along columns, the sign of individual sulcal phenotypes (average depth, depth variability, longest branch, branch span, fractal dimension) used to derive sulcal complexity does not necessarily indicate the sign of sulcal complexity, unlike what is observed for these phenotypes when TTV-corrected. This occurs because sulcal complexity captures relative strengths of individual phenotypes within each brain. For example, in 3q29 deletion syndrome, where microcephaly is observed, constituent sulcal phenotypes TTV-uncorrected effects mostly reflect the residual TTV effect: widespread decreases in longest branch length and fractal dimension across the cortex are consistent with decreased TTV. See **Supplementary Data 1** for all effect sizes and associated *p*-values.

**B)** Distribution of TTV-uncorrected absolute effect sizes across cortical sulci were plotted for each phenotype and for each syndrome. Box plots were ordered for each phenotype by the rank of median strength of sulcal complexity effects. Strength of associations vary from those seen in TTV-corrected sulcal phenotypes due to absolute effects of TTV on individual sulcal phenotypes.

**C)** Spatial correlations were determined between each TTV-uncorrected phenotype's standardized effects and TTV-uncorrected sulcal complexity standardized effects for each syndrome. Significance was corrected for tests across syndrome and adjusted for effects of spatial auto-correlation (Bonferroni: $p_{spin} < 0.05/9$, nominal: $p_{spin} <$

0.05). Despite changes in the cortex-wide mean values for TTV-uncorrected sulcal phenotypes, the relative spatial patterning coheres with sulcal complexity in the same manner as observed with TTV-corrected phenotype effects. Again, average depth and branch span were most linked to the spatial patterning of sulcal complexity, and no significant relationships were observed with sulcal surface area and sulcal thickness.

**D)** Finally, we directly compared the spatial similarity between TTV-corrected and TTV-uncorrected phenotypes. All spatial correlations for any phenotype and syndrome pair were significant (Bonferroni: $p_{spin} < 0.05/9$). Therefore, the relative patterning of sulcal phenotype effects in neurogenetic syndromes is not altered by correction for TTV.

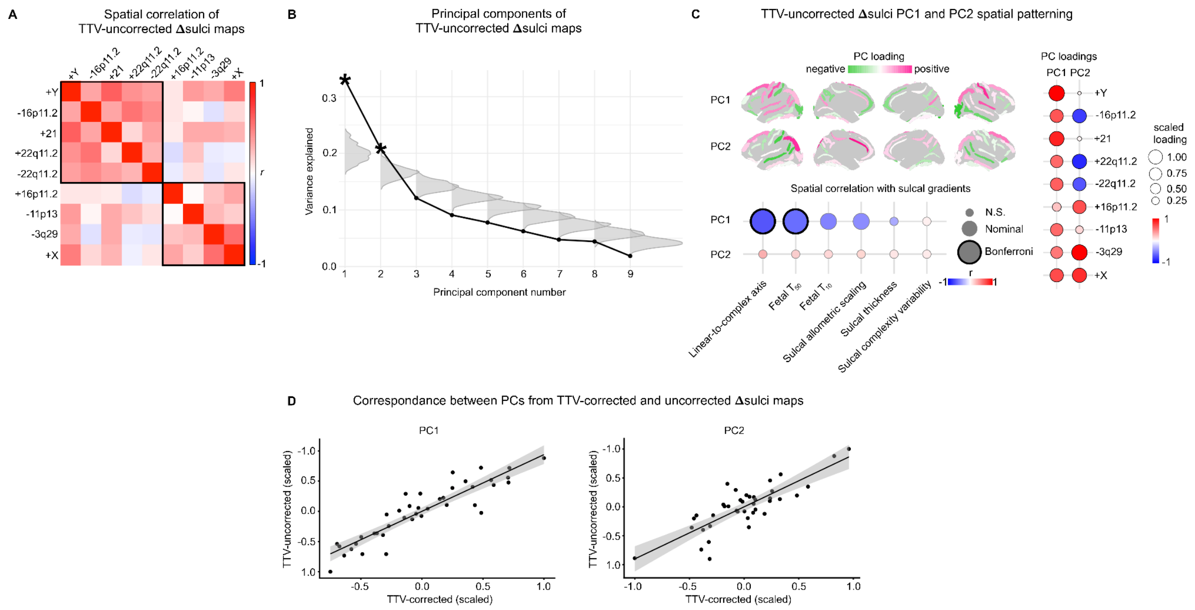


Supplementary Figure 3. **The shared axis of ∆complexity effects is replicable with TTV-uncorrected ∆complexity estimates.**

**A)** Spatial correlations between TTV-uncorrected ∆complexity effect maps are shown, showing similar cross-syndrome spatial correlations and clustering as demonstrated with TTV-corrected ∆complexity.

**B)** Just as with TTV-corrected ∆complexity, the first two PCs of TTV-uncorrected ∆complexity effects explain significantly more variance than when randomly permuting spatial maps (gray distribution) ($p_{perm} < 0.05$).

**C)** TTV-uncorrected ∆complexity PC1 and PC2 are plotted on the cortical surface, with the sign of PC scores chosen to be positively correlated with the syndrome TTV-uncorrected ∆complexity effects that most strongly load on respective PCs. TTV-uncorrected ∆complexity PC1 and PC2 mirror the spatial patterning and relationships with syndromes and sulcal gradients as observed for TTV-corrected ∆complexity. See **Supplementary Data 2** for both TTV-corrected and TTV-uncorrected ∆complexity PC1 and PC2.

**D)** Finally, we directly tested the correspondence between TTV-uncorrected and TTV-uncorrected ∆complexity PCs. PC1 (spatial correlation $r = 0.91$, $p_{spin} < 0.05$) and PC2 (spatial correlation $r = 0.82$, $p_{spin} < 0.05$) were both strongly replicated when using TTV-uncorrected ∆complexity.

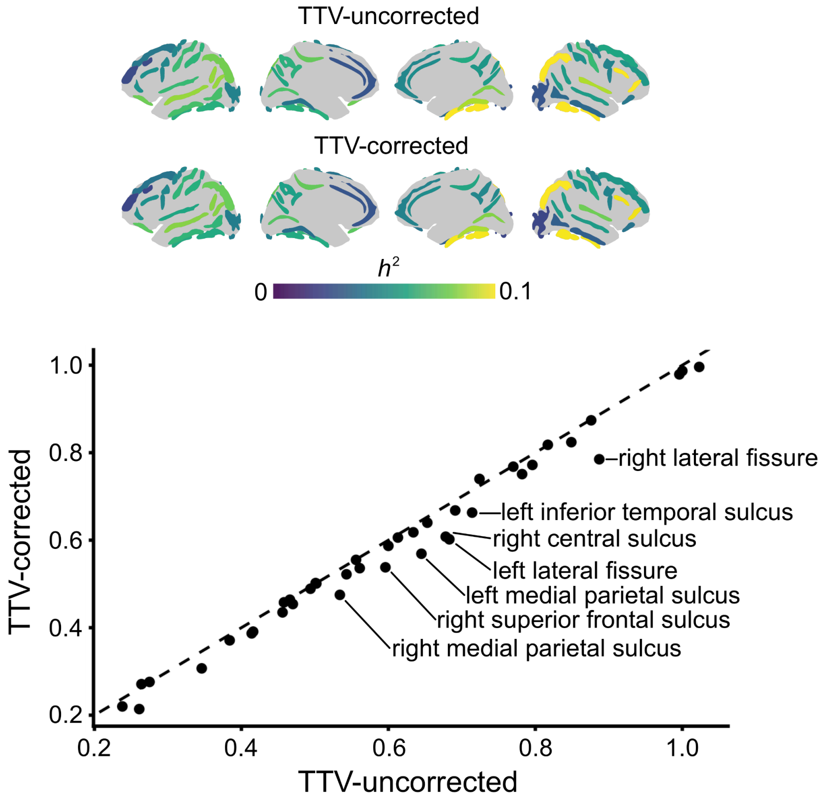


Supplementary Figure 4. **TTV regression effect on sulcal complexity heritability.** SNP-based heritability was computed for sulcal complexity with and without regression of total tissue volume (TTV) of the cortex. **(Top)** The maps are visually similar both in terms of the magnitude of heritability and the inter-regional patterning of its values from sulcus to sulcus, indicating that the genetic architecture of sulcal complexity is not biased by TTV effects. **(Bottom)** The spatial correlation across sulci for SNP-based heritability is shown (r = 0.99), with slope that does not significantly deviate from 1 ($p > 0.05$). Labeled points highlight deviations greater than 0.005 between TTV-corrected and TTV-uncorrected sulcal complexity heritability. A slight right shift of points from the $y$=$x$ line is observed, due to additional heritability being captured from shared TTV volume in TTV-uncorrected sulcal complexity. See **Supplementary Data 3** for TTV-corrected and TTV-uncorrected sulcal complexity SNP-based heritability estimates.

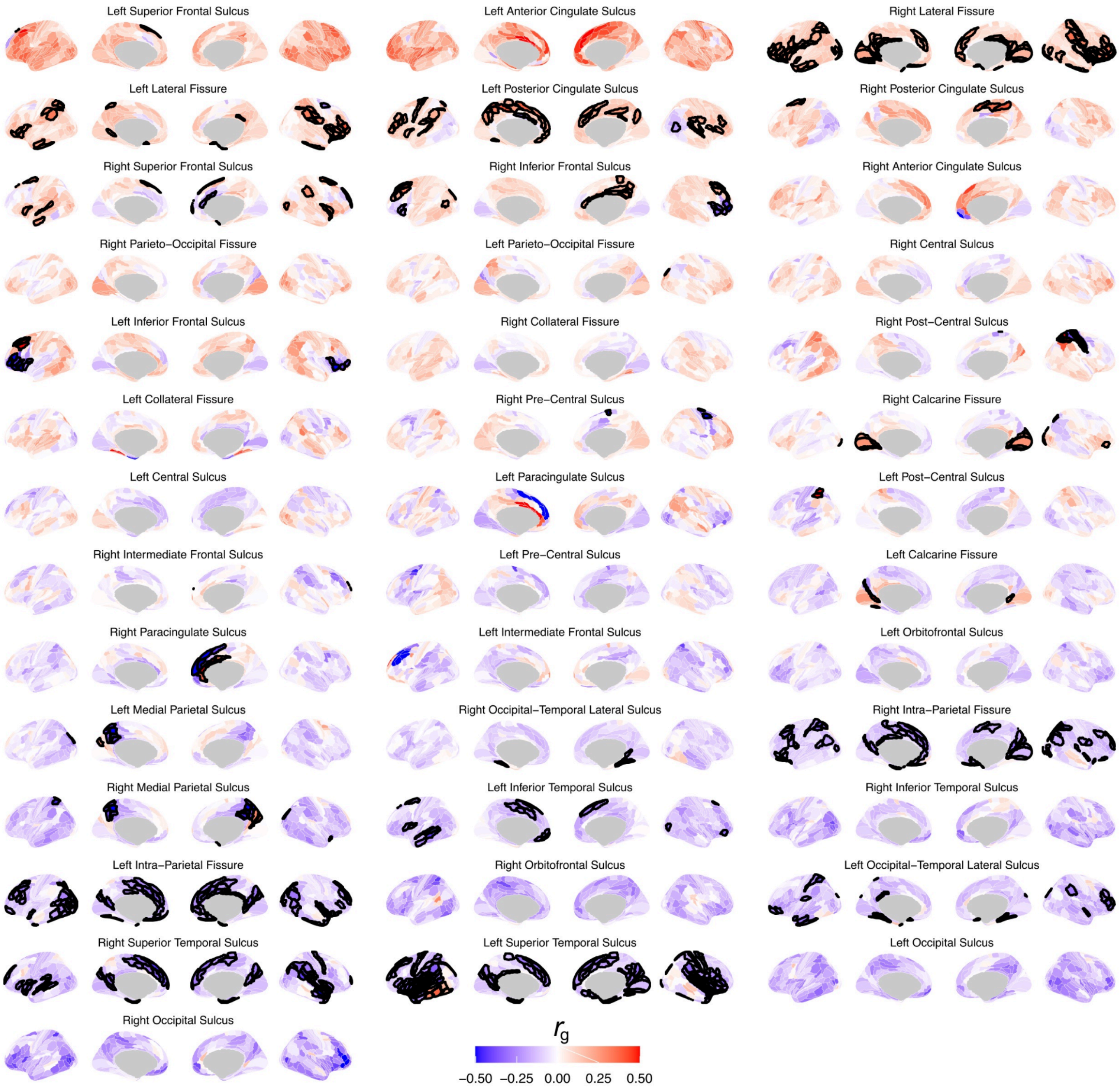


Supplementary Figure 5. **Genetic correlations between sulcal complexity and regional surface area.** Local and global patterns of genetic correlations are observed, with significant positive and negative associations bolded and corrected across cortical parcels (FDR $q$ < 0.05). Sulci are listed in order of their loading on the first principle component of the pairwise sulci x cortical region matrix ({40 sulci sulcal complexity x 360 regions surface area}). This ordering highlights a global pattern whereby genetic variants that impart globally greater cortical surface area appear to reinforce the linear-to-complex axis of sulcal morphology. However, the strongest and most significant associations highlight locally distinct genetic correlations between sulcal complexity and area with the regions surrounding a sulcus. See **Supplementary Data 4** for all genetic correlations and $p$-values.

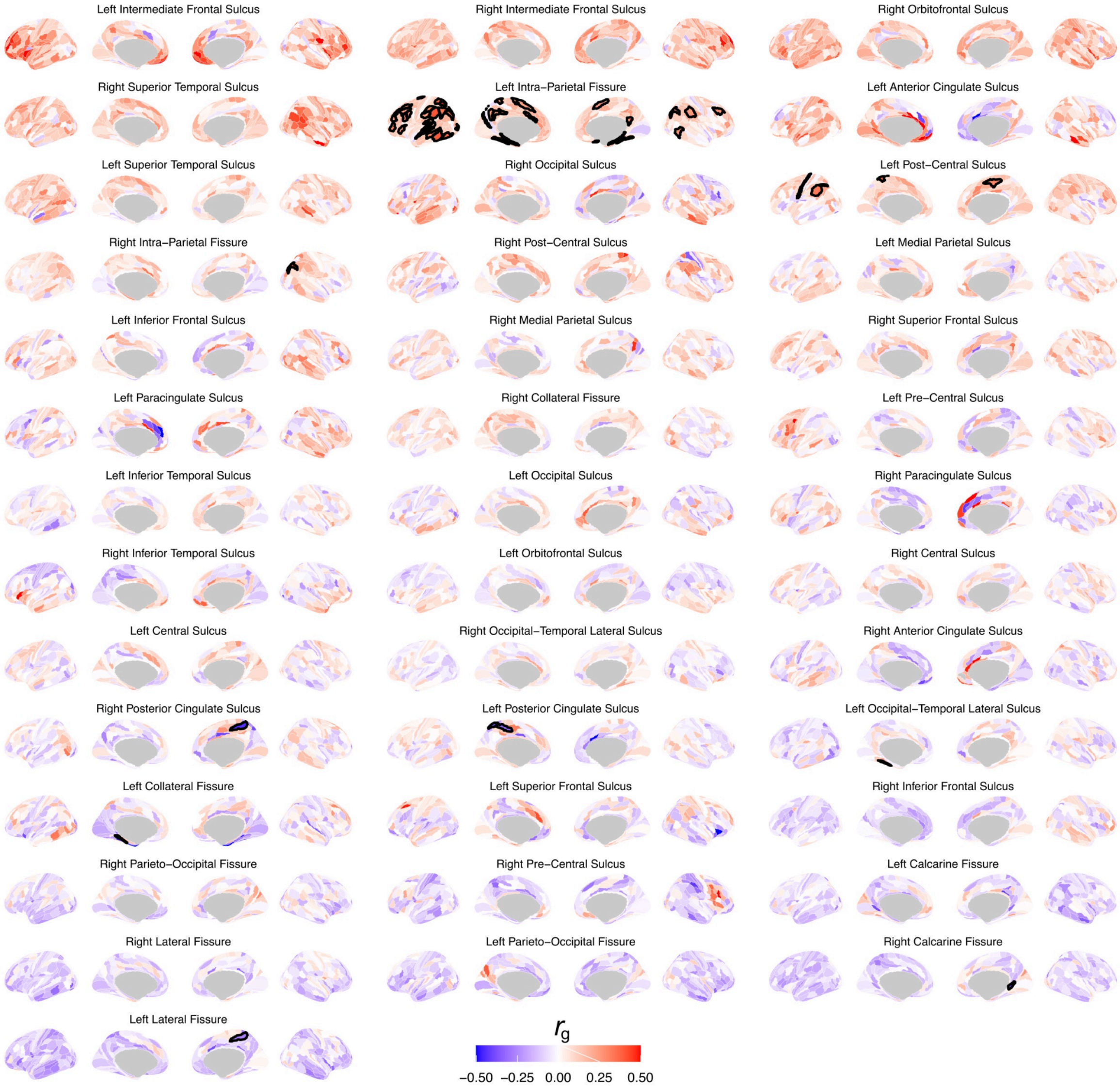


Supplementary Figure 6. **Genetic correlations between sulcal complexity and cortical thickness.** Local and global patterns of genetic correlations are observed, with significant positive and negative associations bolded and corrected across cortical parcels (FDR $q < 0.05$). Sulci are listed in order of their loading on the first principle component of the pairwise sulci x cortical region matrix ({40 sulci sulcal complexity x 360 regions surface thickness}). This ordering highlights a global pattern whereby genetic variants that impart globally decreased cortical surface area appear to reinforce the linear-to-complex axis of sulcal morphology. However, the strongest and most significant associations highlight locally distinct genetic correlations between sulcal complexity and thickness for regions overlapping with a sulcus. See **Supplementary Data 4** for all genetic correlations and *p*-values.

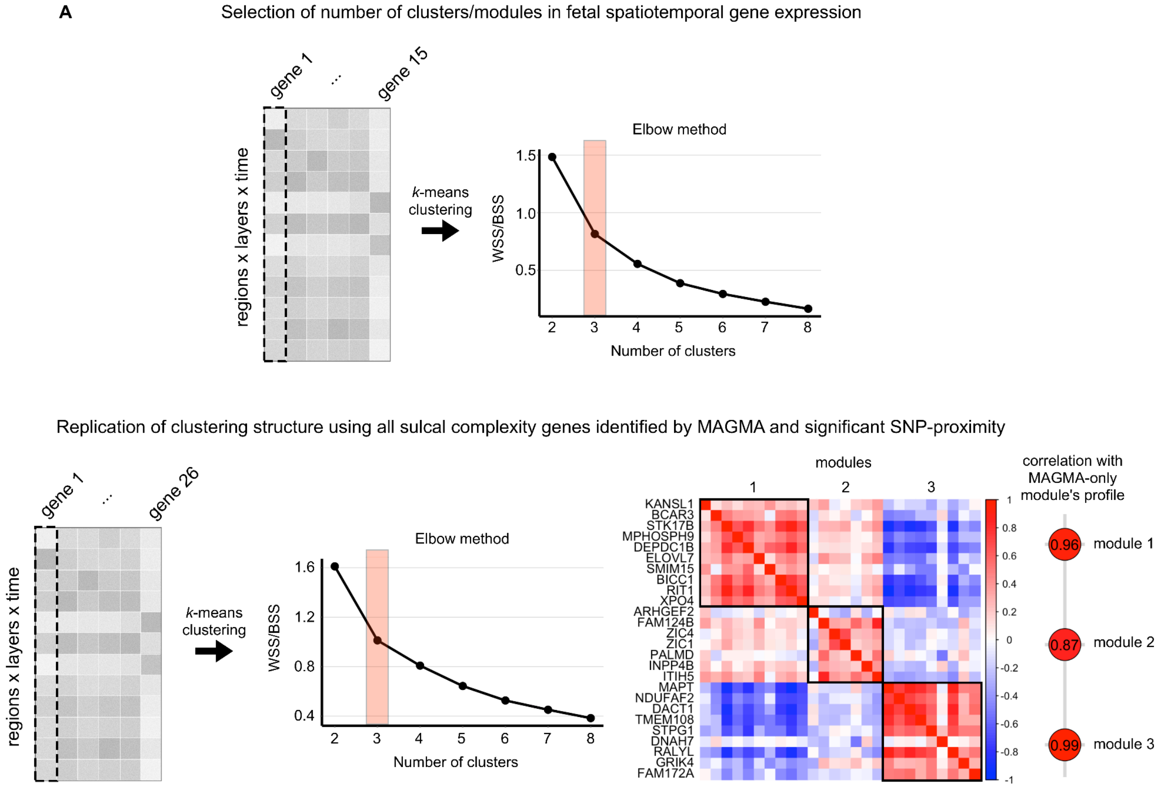


Supplementary Figure 7. ***K*-means clustering and reproducibility of sulcal complexity gene modules in fetal spatiotemporal gene expression data.**

**A)** The k-means clustering used to identify the modules of sulcal complexity genes for the 15 genome-wide MAGMA significant genes for which data was available in the preprocessed µBrain fetal brain expression dataset. Spatiotemporal data across 20 cortical regions, 5 tissue layers, and 2 gestational time points were aggregated for clustering. The elbow method was used to find a set of biologically valid clusters, balancing between over and underfitting. The number of clusters/modules identified was determined from the elbow in a plot of clusters/modules versus the ratio of within sum of squared differences and between sum of squared distances (WSS/BSS) across clusters. $k = 3$ was chosen as decreases in WSS/BSS are significantly attenuated for subsequent values of $k$.

**B)** The same approach for clustering was repeated, but instead using the set of all 50 candidate sulcal complexity genes for which data was available in the µBrain dataset. These 26 genes had similar clustering properties, both by having an optimal $k = 3$ and by yielding similar groupings of genes in clusters/modules as observed for the 15 genome-wide MAGMA significant genes. The degree-weighted average spatiotemporal expression was computed for each module and correlated with the corresponding modules identified from the 15 genome-wide MAGMA significant genes. High correlations demonstrate that different subsets of sulcal complexity genes stably identify the same three modules of fetal brain gene expression.

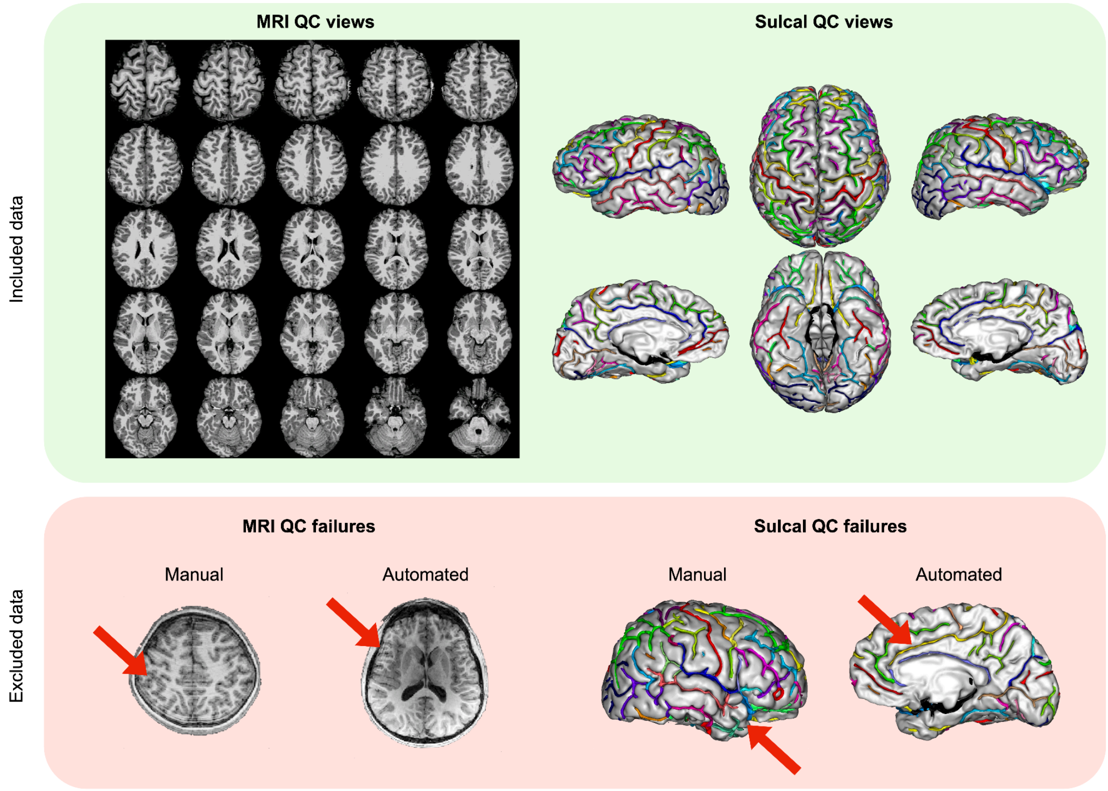


Supplementary Figure 8. **Quality control (QC) of imaging data from neurogenetic syndrome cohorts.**
**(Top)** Data that passed the four quality control measures are displayed with the image layout generated for manual MRI and sulcal QC. For manual MRI QC, head motion related artifacts are dominantly visible in the axial plane, so axial slices were generated from superior to inferior directions for each subject. For manual sulcal QC, 6 views covered all angles for which sulci populate the cortical sheet. These images generated for each subject and were inspected for at least one gross abnormality in sulcal reconstruction that did not reflect the sulcal paths in underlying MRI data.
**(Bottom)** Data that did not pass quality control in 4 example subjects. Manual MRI QC failure was due to identification of ringing artifacts that obscured gray-white tissue boundaries and sulcal paths. Automated MRI QC using Euler number (EN) used the threshold EN > -217 for passing data. The example subject had substantially lower EN (EN = -1078), which indexed MRI quality as seen in the significant blurring artifact in the image. Importantly, automated QC alone would not have been sufficient as the example for manual MRI QC had EN = -196. For manual sulcal QC, the gray and white matter segmentation failed to accurately represent the temporal pole region, leading to unreliable sulcal reconstruction and sulcal measurement. The automated sulcal QC involved checking whether all 40 sulci were labeled by BrainVISA Morphologist's automated sulcal labeling (Borne et al., 2020). This subject lacked a label for the anterior cingulate sulcus even though it is clearly present. The posterior cingulate sulcus (yellow) absorbed the anterior cingulate sulcus' label as the sulcal segments were failed to be automatically split into anterior and posterior sulcal regions. Such mislabeling behavior can happen between any pair of adjacent sulci, necessitating this step in sulcal QC.

| Cohort | Group | *n* (% female) | Age, mean (SD) [range] | EN, mean (SD) |
|---|---|---|---|---|
| +X | control | 54 | 18.0 (5.4) [6.0-25.0] | -132.1 (35.5) |
| | case | 74 | 16.9 (4.3) [8.1-25.6] | -121.7 (38.2) |
| +Y | control | 35 | 15.1 (4.6) [5.6-24.7] | -122.3 (37.0) |
| | case | 30 | 15.2 (6.1) [6.1-25.9] | -131.7 (39.4) |
| -3q29 | control | 54 (42.6) | 18.1 (9.0) [4.8-40.8] | -100.3 (41.0) |
| | case | 14 (42.9) | 18.5 (9.4) [4.9-39.1] | -104.9 (45.7) |
| -11p13 | control | 19 (57.9) | 18.1 (8.8) [7.9-34.9] | -99.2 (42.4) |
| | case | 31 (45.2) | 14.9 (8.6) [6.1-33.4] | -110.6 (43.8) |
| -16p11.2 | control | 23 (52.2) | 30.9 (11.9) [7.8-52.1] | -91.7 (42.2) |
| | case | 16 (50.0) | 24.8 (12.8) [9.2-53.6] | -94.6 (32.5) |
| +16p11.2 | control | 23 (43.5) | 37.1 (13.2) [9.5-62.5] | -93.4 (39.5) |
| | case | 20 (45.0) | 32.4 (12.7) [9.8-58.1] | -103.5 (43.9) |
| +21 | control | 40 (50.0) | 16.2 (5.6) [6.4-24.3] | -115.1 (45.8) |
| | case | 22 (50.0) | 16.4 (5.7) [6.8-24.5] | -90.9 (41.0) |
| -22q11.2 | control | 32 (53.1) | 14.2 (5.6) [6.0-29.1] | -97.0 (45.0) |
| | case | 72 (51.4) | 14.9 (6.2) [5.6-34.4] | -104.4 (42.9) |
| +22q11.2 | control | 32 (53.1) | 13.4 (6.4) [6.1-28.9] | -108.3 (45.4) |
| | case | 24 (41.7) | 12.9 (5.8) [7.4-32.3] | -124.4 (39.6) |

Supplementary Table 1. **Demographics of each neurogenetic syndrome cohort.** Case and control groups within each neurogenetic syndrome cohort were matched on age, sex, site, and EN. Lower EN indicates poorer scan quality.

| Cohort | Ascertainment and inclusion | Scanning protocol | References |
|---|---|---|---|
| XXY, XYY | All participants had normal radiological reports and no prior brain injuries. XXY or XYY diagnosis was confirmed by karyotype, all patients non-mosaic. All control participants were screened using a structured interview to exclude a history of neurodevelopmental or psychiatric disorders. | Scanner: MR750 3T scanner (General Electric)<br>Sequence: T1-weighted magnetization-prepared rapid acquisition gradient-echo (MPRAGE) with (172 contiguous sagittal slices with 256 x 256 in-plane matrix and 1 mm slice thickness yielding 1 mm isotropic voxels) | Levitis et al. (2024) |
| 3q29 deletion | 3q29Del participants were ascertained from the 3q29 Registry. 3q29Del status was confirmed via clinical genetics reports and/or medical records. | Scanner: 3T Siemens Magnetom Prisma scanner, using a 32-channel head coil and an 80mT/m gradient<br>Sequence: T1-weighted MPRAGE were acquired in the sagittal plane using a single-echo sequence with the following parameters: TE = 2.24 ms, TR = 2400 ms, TI = 1000 ms, bandwidth = 210 Hz/pixel, FOV = 256x256mm, Resolution = 0.8 mm isotropic | Sefik et al. (2024) |
| 11p13 deletion | Chromosome deletions were characterized by microsatellite marker analysis and oligonucleotide array comparative genomic hybridization. Healthy controls were screened and excluded for history of neurological and psychological impairments. | Scanner: 3.0 T Philips Achieva MRI scanner equipped with an 8-channel phased array head coil<br>Sequence: 3D TFE T1-weighted sequence with scan parameters: TR = 8.3 ms, TE = 3.8 ms, TI delay = 1031 ms, 160 shots. In total, 171 slices were acquired in the sagittal plane with an acquisition matrix of 240 × 240 and an FOV of 240 mm | Han et al. (2013)<br>Seidlitz et al. (2020) |
| 16p11.2 deletion and duplication | In the European 16p11.2 consortium, the families were directly recruited by the referring physician and had various genetic tests to identify the CNV. Controls were either non-carriers within the same families or individuals from the general population. | Scanner: Two 3T scanners were used, either a Magnetom TIM Trio (Siemens Healthcare, Erlangen, Germany), using a 12-channel RF receive head coil and RF body transmit coil or a Magnetom Prisma Syngo (Siemens Healthcare, Erlangen, Germany) using a 64-channel RF receive head coil and RF body transmit coil.<br>Sequence: On the TIM Trio scanner, T1-weighted (T1w) anatomical images acquired using a Multi-Echo Magnetization Prepared RApid Gradient Echo sequence (ME-MPRAGE: 176 slices; 256×256 matrix; echo time (TE): TE1 = 1.64 ms, TE2 = 3.5 ms, TE3 = 5.36 ms, TE4 = 7.22 ms; repetition time (TR): 2530 ms; flip angle 7°). On the Prisma Syngo scanner, T1w images were acquired using a single-echo MPRAGE sequence (176 slices; 256×256 matrix; TE = 2.39 ms; TR = 2000 ms; flip angle 9°). Consistent with recent studies (Kumar et al., 2025; Modenato et al., 2021) and in line with negligible site effects observed in this cohort (Martin-Brevet et al., 2018), we chose not to explore further site effects in this cohort in our analyses. | Modenato et al. (2021)<br>Martin-Brevet et al. (2018)<br>Jacquemont et al. (2011)<br>Kumar et al. (2025) |
| Trisomy 21 | All participants had normal radiological reports and no prior brain injuries. All participants with trisomy 21 had chromosomal diagnoses of trisomy 21 according to genetic testing or parent report. All participants with trisomy 21 with follow-up genetic testing were non-mosaic. All control participants were screened using a structured interview to exclude a history of neurodevelopmental or psychiatric disorders. | Scanner: 3T General Electric Scanner using an 8-channel head coil.<br>Sequence: High-resolution (0.94 × 0.94 × 1.2 mm) T1-weighted images were acquired utilizing an ASSET-calibrated magnetization prepared rapid gradient echo sequence (128 slices; 224 × 224 acquisition matrix; flip angle = 12°; field of view [FOV] = 240 mm). | Lee et al. (2016)<br>Levitis et al. (2024) |
| 22q11.2 deletion and duplication | All 22q11.2 CNV carriers had molecularly confirmed 22q11.2 deletions or duplications. Exclusion criteria for all study participants included significant neurological or medical conditions (unrelated to 22q11.2 CNV) that might affect brain structure, history of head injury with loss of consciousness, insufficient fluency in English, and/or substance or alcohol abuse or dependence within the past 6 months. TD controls could not have significant intellectual disability or meet criteria for any major mental disorder. | Scanner: Two 3T scanners were used, either a Siemens Tim Trio MRI scanner (12-channel head coil), or a Siemens Prisma scanner (32-channel head coil).<br>Sequence: The parameters for the MPRAGE on the Siemens Tim Trio were the following: TR = 2.3 s, TE = 2.91 ms, FOV = 256 mm, matrix = 240 × 256, flip angle = 9°, slice thickness = 1.20 mm, 160 slices. Prisma MPRAGE acquisition parameters were almost identical: TR = 2.3 s, TE = 2.94 ms, FOV = 256 mm, matrix = 240 × 256, flip angle = 9°, slice thickness = 1.20 mm, 160 slices. | Jalbrzikowski et al. (2022) |
| UK Biobank | All subjects were typically developed adults without neurological conditions. | Scanner: All three scanning sites used standard Siemens Skyra 3T scanners with a Siemens 32-channel RF receive head coil<br>Sequence: T1 acquisition involved a five-minute 3D MPRAGE session at 1x1x1 mm resolution, in-plane acceleration iPAT=2, and prescan-normalization. T2-FLAIR acquisition involved a six-minute 3D SPACE session at 1.05x1x1 mm resolution, in-plane acceleration iPAT=2, partial Fourier = 7/8, fat saturation, elliptical k-space scanning, and pre-scan normalization. | Alfaro-Almagro et al. (2018)<br>Snyder et al. (2024)<br>See also:<br>https://biobank.ctsu.ox.ac.uk/crystal/crystal/docs/brain_mri.pdf |

Supplementary Table 2. **Additional cohort information on ascertainment, inclusion criteria, and scanning protocols.**

**Description of Additional Supplementary Data Files**

Supplementary Data 1. Rare neurogenetic syndrome standardized effect sizes and *p*-values associated with sulcal complexity, average depth, depth variability, longest branch, branch span, fractal dimension, sulcal area, and sulcal thickness – with values given from linear models both with and without correction for total brain tissue volume.

Supplementary Data 2. The two significant principal components of atypical sulcal complexity across rare neurogenetic syndromes (∆complexity PC1 and PC2), both with and without correction for total brain tissue volume.

Supplementary Data 3. SNP-based heritability of sulcal complexity for 40 cortical sulci, with and without correction for total brain tissue volume.

Supplementary Data 4. Genetic correlations between sulcal complexity and cortical area and thickness across 360 cortical parcels on the Glasser atlas.

Supplementary Data 5. Summaries of sulcal complexity genes, their genomic locations, functions, and how they were identified in analyses – MAGMA and SNP-proximity based approaches were used to identify candidate sulcal complexity genes, using both experiment (strict; MAGMA: $p < 7.62 \times 10^{-8}$, SNP: $p < 1.47 \times 10^{-9}$) and genome-wide (relaxed; MAGMA: $p < 2.59 \times 10^{-6}$, SNP: $p < 5 \times 10^{-8}$) thresholds.

Supplementary Data 6. Three modules of spatiotemporal gene expression in the fetal brain across 20 cortical regions, 5 tissue layers, and two timepoints.

Supplementary Data 7. Gene set enrichment analysis results of ranked gene lists produced from spatiotemporal correlation with the three identified modules of sulcal complexity gene expression.
